# Hotspots of aberrant enhancer activity in fibrolamellar carcinoma reveal molecular mechanisms of oncogenesis and intrinsic drug resistance

**DOI:** 10.1101/2020.01.18.911297

**Authors:** Timothy A. Dinh, Ramja Sritharan, F. Donelson Smith, Adam B. Francisco, Rosanna K. Ma, Rodica P. Bunaciu, Matt Kanke, Charles G. Danko, Andrew P. Massa, John D. Scott, Praveen Sethupathy

## Abstract

Fibrolamellar carcinoma (FLC) is a rare, therapeutically intractable liver cancer that disproportionately affects youth. Although FLC tumors exhibit a distinct gene expression profile, the causative transcriptional mechanisms remain unclear. Here we used chromatin run-on sequencing to discover approximately 7,000 enhancers and 141 enhancer hotspots activated in FLC relative to non-malignant liver. Detailed bioinformatic analyses revealed aberrant ERK/MEK signaling and candidate master transcriptional regulators. We also defined the genes most strongly associated with aberrant FLC enhancer activity, including *CA12* and *SLC16A14*. Treatment of FLC cell models with inhibitors of CA12 or SLC16A14 independently reduced cell viability and/or significantly enhanced the effect of MEK inhibitor cobimetinib. These findings highlight new molecular targets for drug development as well as novel drug combination approaches.

## Introduction

Fibrolamellar carcinoma (FLC) is a rare type of liver cancer that predominantly affects adolescents and young adults with no prior history of liver disease (Craig et al., 1980; Torbenson, 2012). Currently, surgical resection is the only effective treatment for FLC, however most patients have metastatic disease at the time of diagnosis, making surgical cures difficult (Stipa et al., 2006). While some patients have been successfully treated with chemotherapy and molecular therapies, there is no standard treatment regimen for FLC patients (Torbenson, 2012). Furthermore, FLC is often drug resistant and frequently recurs following initial treatment (Maniaci et al., 2009), underscoring the need to develop effective therapies for this cancer.

FLC is genetically characterized by a ∼400 kb heterozygous deletion on chromosome 19 that leads to the formation of the *DNAJB1-PRKACA* fusion (Honeyman et al., 2014). This fusion occurs in at least 80% of patients (Cornella et al., 2014; Honeyman et al., 2014), is specific to FLC (Dinh et al., 2017; Graham et al., 2015; Kastenhuber et al., 2017), and is sufficient to drive liver tumor formation in mice (Engelholm et al., 2017; Kastenhuber et al., 2017). Multiple groups have performed genome-scale analyses to identify dysregulated genes (Cornella et al., 2014; Dinh et al., 2017; Griffith et al., 2016; Malouf et al., 2014; Simon et al., 2015; Sorenson et al., 2017; Xu et al., 2014), long non-coding RNAs (Dinh et al., 2017), and microRNAs (Dinh et al., 2019; Farber et al., 2018) in FLC. Yet, little is known about the causative transcriptional regulatory mechanisms that lead to aberrant gene expression and FLC tumor formation.

Precise spatial and temporal regulation of gene expression is essential to many biological processes. One class of *cis*-regulatory elements that plays a major role in transcriptional regulation of gene expression is enhancers. Enhancers are classically defined as stretches of non-coding DNA that promote transcription of target gene(s) irrespective of genomic context, orientation, and, to a substantial extent, distance as well (Blackwood and Kadonaga, 1998). Enhancers are often cell-type specific, allowing precise spatiotemporal control of gene transcription in different cell types within an organism (Heintzman et al., 2009; Nord et al., 2013). Recent work suggests that there are at least tens of thousands of active enhancers in any given cell type (Dunham et al., 2012). Active enhancers serve as binding sites for transcription factors, transcriptional coactivators, and RNA polymerase and are thought to interact with their cognate promoters through three-dimensional looping, explaining their ability to act over long distances (Long et al., 2016).

Recent studies have shown that active enhancers are transcribed to produce enhancer RNAs (eRNAs; Kim et al., 2010; De Santa et al., 2010). Genome-wide identification of enhancers by detection of eRNAs has recently become possible by coupling nascent transcription (run-on) assays with high throughput sequencing (e.g., global run-on [GRO-seq] and precision run-on sequencing [PRO-seq]). GRO-seq (Core et al., 2008) and PRO-seq (Kwak et al., 2013) require isolation of cellular nuclei making the application of these methods to primary tissue extraordinarily difficult. To overcome this limitation, chromatin run-on sequencing (ChRO-seq), was developed to extend the technique and permit analysis of primary fresh or frozen tissue (Chu et al., 2018). Additionally, the advent of a sister technique called length extension ChRO-seq (leChRO-seq) permits investigation of samples with degraded RNA (Chu et al., 2018). These technical advances finally enable the study of nascent transcription at enhancer, promoter, and gene loci in fresh or archived primary human tumor tissues. ChRO-seq was recently successfully used to identify distinct transcriptional programs in different subtypes of glioblastoma multiforme (Chu et al., 2018).

Here, we perform ChRO-seq in primary FLC and matched non-malignant liver (NML) samples. As FLC is a rare adolescent cancer, we worked closely with the Fibrolamellar Cancer Foundation (FCF; https://fibrofoundation.org) over multiple years to accumulate the samples for this study. In order to bolster the number of NML samples, we also leverage publicly available enhancer and super enhancer data generated for an additional 14 human liver-derived samples from the super enhancer database SEdb (Jiang et al., 2019). By integrating our ChRO-seq analyses with RNA-seq data, we identified 16 genes strongly associated with aberrant enhancer activity in FLC. Overall, our study defines for the first time the unique enhancer activity profile and master transcription factors in FLC, identifies high-confidence candidate targets of the perturbed transcriptional regulatory programs, and indicates that MAPK inhibition, as well as SLC16A14 or CA12 inhibition alone or in combination, represent molecular therapeutic strategies to study further in FLC.

## Results

### Mapping nascent transcription and transcriptional regulatory elements in FLC

To identify transcriptional regulatory elements (TREs) in FLC, we performed (le)ChRO-seq on 14 FLC samples and 3 matched NMLs (Fig. 1A, Tables S1,S2). All FLC tumor samples expressed the DNAJB1-PRKACA fusion (Table S2). The bioinformatic pipeline for sequencing data processing and genome mapping is provided in Materials and Methods. After removing adapters and PCR duplicates, we obtained an average of ∼17.9 million mapped reads per sample (Table S2). We observed a buildup of transcriptional signal at transcription start sites (TSS) and the start of exons (Fig. S1A), consistent with previous reports (Core et al., 2008; Kwak et al., 2013). A total of 153,478 unique TREs across all samples were detected using dREG, a previously published algorithm that identifies transcriptional regulatory elements from run-on sequencing experiments (Danko et al., 2015; Wang et al., 2019). The majority of TREs are embedded in intronic (43.98%) and intergenic (31.49%) regions with 17.63% of TREs located close to TSSs (Fig. 1B). As expected, TREs close to TSSs are responsible for the majority of the ChRO-seq signal across all TREs (Fig. S1B,C). On average, the TREs are 413 bp in length (Fig. 1C), consistent with the 50-1500 bp length previously proposed for enhancers (Blackwood and Kadonaga, 1998; Parker et al., 2013). We did not observe major differences in TRE length based on genomic context or overlap with CpG islands (Fig. S1D,E). Hierarchical clustering and principal component analysis demonstrate that transcription at TREs stratifies FLC from NML samples (Fig. 1D,E). Interestingly, clustering based on transcription at distal TREs maintains the stratification between FLC and NML, whereas clustering with proximal TREs does not (Fig. 1D,E). Distal TREs (hereafter, referred to as enhancers) stratify FLC and NML better than gene body transcription, mRNA expression profiles, and microRNA expression profiles (Fig. 1D,E; Fig. S1F), indicating that enhancer activity is more cell-type and condition-specific than these other data types.

**Fig. 1.**
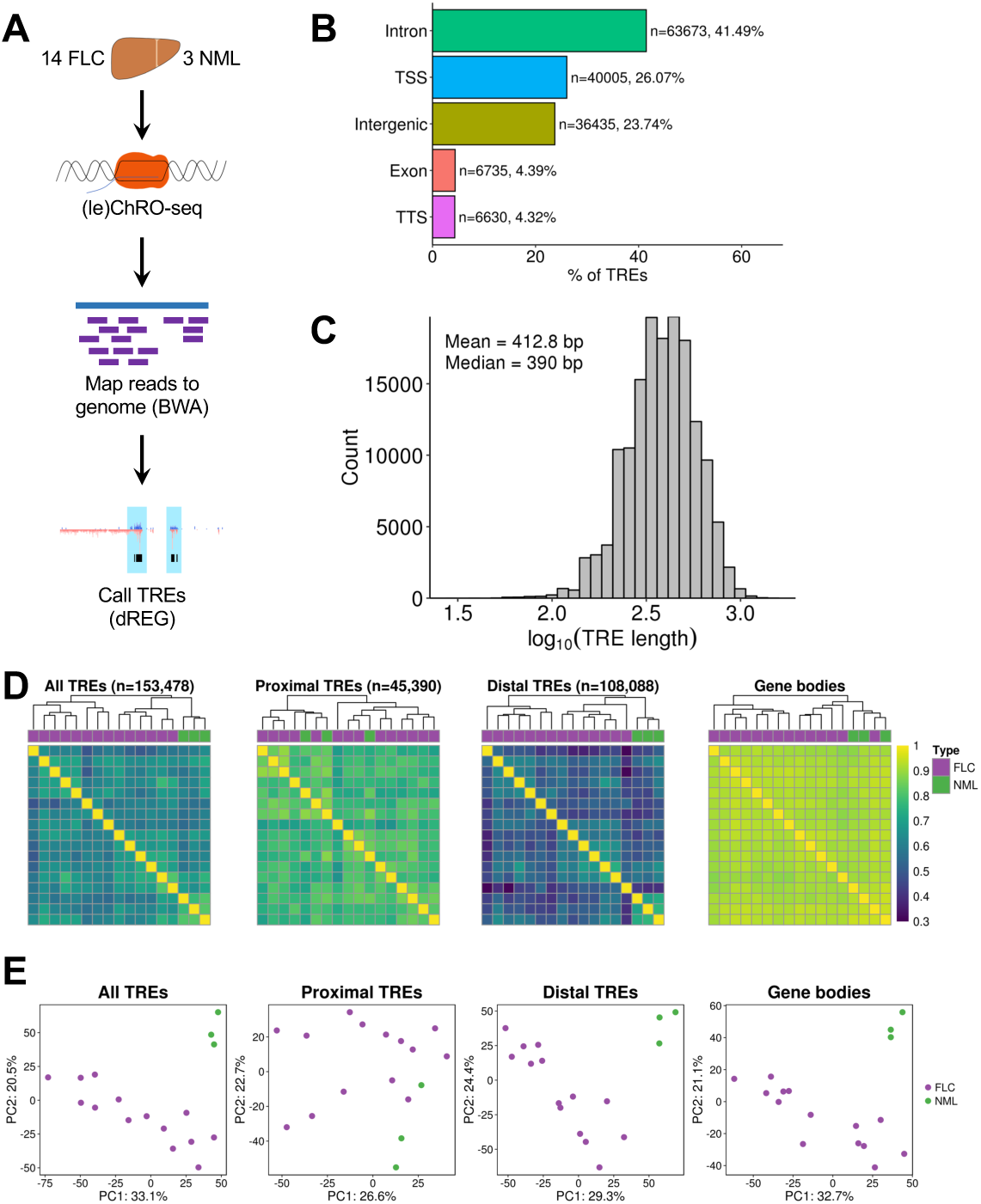
ChRO-seq reveals TREs that stratify FLC from NML. (A) Diagram of ChRO-seq workflow. (B) Bargraph showing the distribution of genomic locations of TREs identified by ChRO-seq. Genomic locations are defined by HOMER using GENCODE v25 annotations. (C) Length distribution of TREs identified by ChRO-seq. (D) Heatmap of pairwise correlations from TRE transcription. Hierarchical clustering was performed using Euclidean distance and Ward’s minimum variance method. Color bar shows Spearman’s correlation coefficient. TREs were classified based on GENCODE v25 basic annotations. (E) Principal components analysis of TRE transcription. Analyses were performed using the 1000 most variable TREs following variance stabilizing transformation (DESeq2). The axes display the first two principle components and the variance explained by each component. PC, principal component; TSS, transcription start site; TTS, transcription termination site; UTR, untranslated region.

### Identification of FLC-specific TREs

In order to identify TREs that are more actively transcribed in FLC relative to NML, we quantified ChRO-seq reads within each TRE across all samples and used the DESeq2 algorithm (Love et al., 2014) to perform differential transcription analysis (Fig. 2A). This approach led to the identification of 6824 TREs that are significantly more actively transcribed in FLC (Fig. 2B,C; FDR < 0.05, log_2_(fold change) > 0). We refer to these as FLC-specific TREs. We also identified 1317 NML-specific TREs. Most FLC- and NML-specific TREs are located in intronic, intergenic, and TSS regions and do not show major differences in TRE length (Fig S2A,B).

**Fig. 2.**
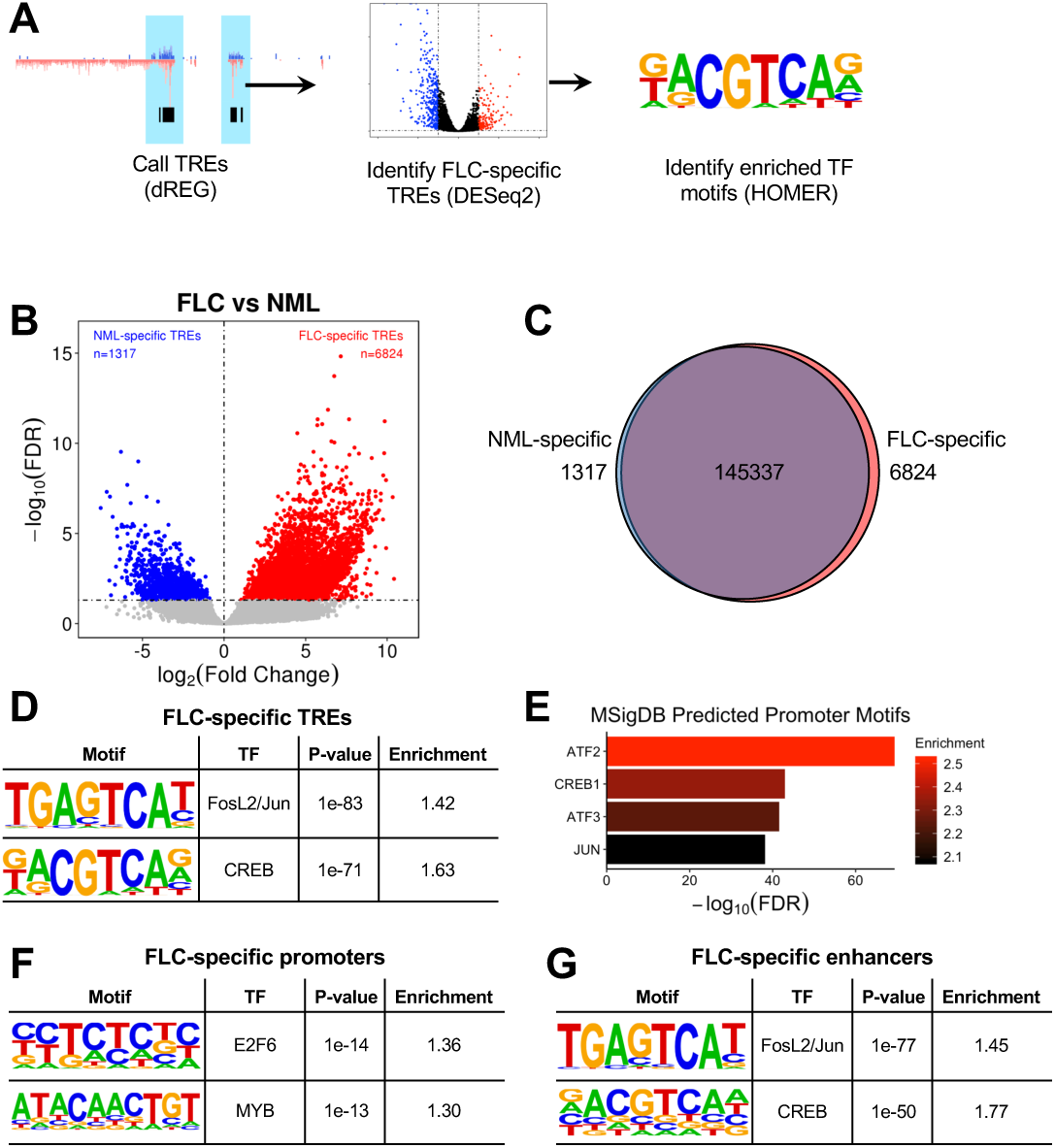
FLC-specific TREs are enriched for FOS/JUN and CREB motifs. **(A)** Diagram of ChRO-seq workflow to identify FLC-specific TREs and enriched transcription factor motifs. **(B)** Volcano plot displaying differentially transcribed TREs in FLC compared to NML. Dashed lines represent FDR = 0.05 (horizontal) and log_2_(fold change) = 0 (vertical). **(C)** Venn diagram showing the total number of TREs and TREs that are FLC- or NML-specific. **(D,F,G)** Tables showing results of transcription factor motif enrichment in FLC-specific TREs compared to non-FLC-specific TREs (D), FLC-specific promoters compared to non-FLC-specific promoters (F), and FLC-specific enhancers compared to non-FLC-specific enhancers (G). Full results are shown in Table S3. **(E)** GREAT analysis of predicted promoter motifs in genes associated with FLC-specific TREs.

Transcription factor motif enrichment analysis of condition-specific TREs using HOMER (Heinz et al., 2010) revealed significant enrichment in FLC-specific TREs for motifs of FOSL2/JUN (AP-1) and CREB (Fig. 2D), both of which are activated by MAPK signaling. Similar analysis of NML-specific TREs showed enrichment for HNF4A motifs (Table S3). Additionally, analysis of the promoters of genes nearest to FLC-specific TREs using the Genomic Regions Enrichment of Annotations Tool (GREAT, McLean et al., 2010) revealed the most significant enrichment for motifs of CREB family members ATF2, CREB1, and ATF3, as well as JUN (Fig. 2E). Separate examination of FLC-specific promoters (proximal TREs) and enhancers (distal TREs) revealed promoter enrichment of E2F6, MYB, and FOSL1 (Fig. 2F, Table S3) and enhancer enrichment of FOSL2/JUN and CREB (Fig. 2G). We found that 2129 (38.2%) FLC-specific enhancers contain FOSL2/JUN motifs and 1090 (19.6%) contain CREB motifs. FLC-specific TREs that did not contain FOSL2/JUN or CREB motifs were enriched in HES2 and SOX1 motifs (Table S3). Our results suggest that FLC is characterized by an aberrant map of regulatory elements that is strongly associated with transcription factors from the CREB, AP-1 families, E2F, and MYB families.

### Identification of genes associated with the highest FLC-specific enhancer density

In order to determine which genes are regulated by FLC-specific TREs, we first defined a genomic window around each TSS. As most promoter-enhancer interactions occur within a few hundred kilobases (Javierre et al., 2016), we defined a 100 kb window upstream and downstream of each gene’s TSS. Next, we quantified the number of FLC-specific enhancers within each window (Fig. 3A). As anticipated, we observed that genes with more FLC-specific enhancers in their windows are more highly transcribed in FLC than NML compared to genes with fewer FLC-specific enhancers in their windows (Fig. 3B). However, it is important to note that only some of the enhancers within a gene’s window are likely to regulate that gene. We address this point more quantitatively later in the manuscript.

**Fig. 3.**
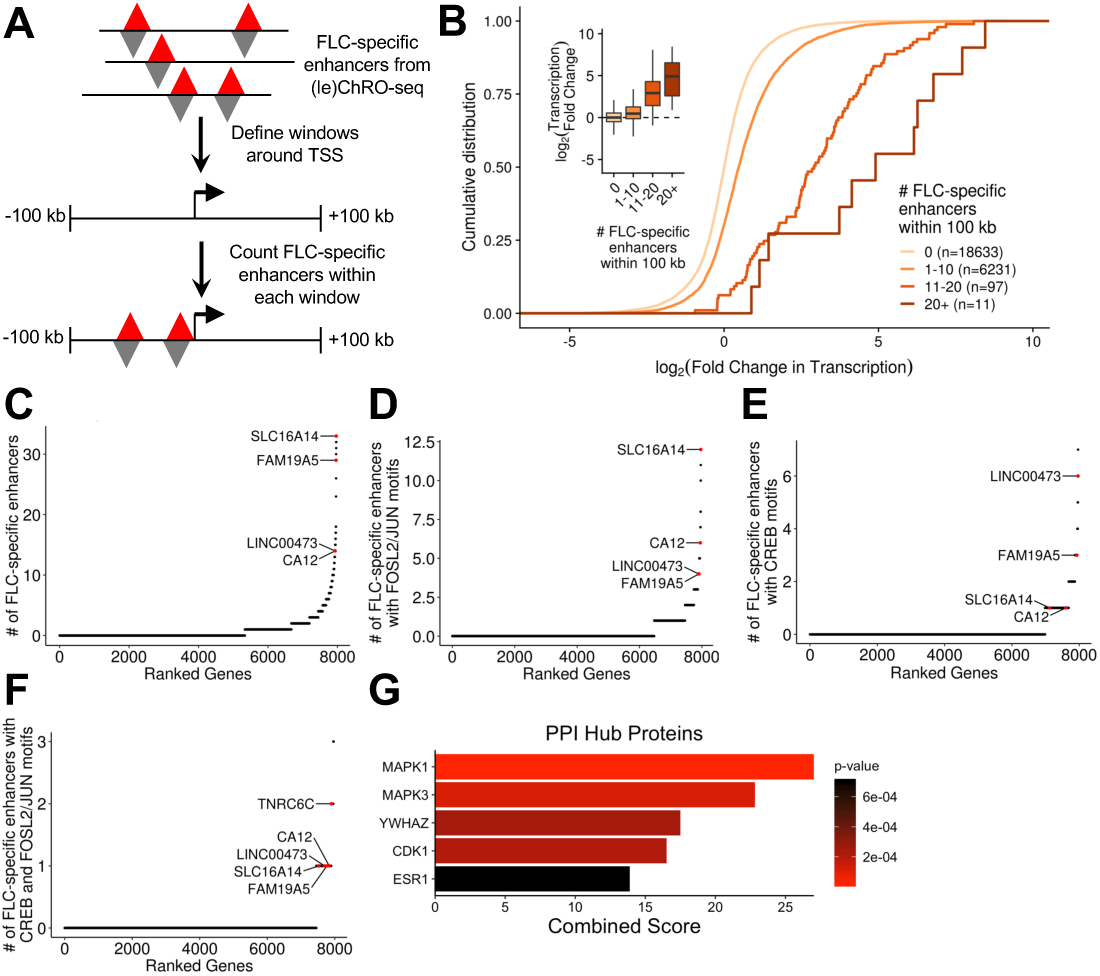
Identification of clusters of FLC-specific enhancers and candidate target genes. **(A)** Schematic of the approach used to link FLC-specific enhancers to candidate genes. Gene windows were defined as 100 kb upstream and 100 kb downstream of each TSS. **(B)** Cumulative distribution function and boxplots (inset) showing the relationship between the number of FLC-specific enhancers within each gene window and the transcriptional fold change in FLC compared to NML. Genes were binned based on the number of FLC-specific enhancers within their gene window and included in the analysis if they were transcribed with a threshold of TPM > 1 in either FLC or NML. **(C,D,E,F)** Genes ranked based on the density of FLC-specific enhancers (C), FLC-specific enhancers with FOSL2/JUN motifs (D), FLC-specific enhancers with CREB motifs (E), or FLC-specific enhancers with both FOSL2/JUN and CREB motifs (F) within their gene windows. Genes were included in the analysis if they were highly expressed (TPM ≥ 25). **(G)** Protein-protein interaction (PPI) hub enrichment of genes with at least one FLC-specific enhancer containing both FOSL2/JUN and CREB motifs within 100 kb of the TSS.

We next ranked genes actively transcribed in FLC (ChRO-seq: TPM ≥ 25) according to the number of FLC-specific enhancers in their windows. The top ranked gene from this analysis is *SLC16A14* (Fig. 3C, Table S4). We also observed that *CA12* and *LINC00473*, two genes that are overexpressed in FLC (Dinh et al., 2017), have very high densities of FLC-specific enhancers. Since we had discovered that motifs of both FOSL2/JUN and CREB are enriched in FLC-specific TREs (Fig. 2D), we next focused on target genes of these transcription factors in FLC. Specifically, we ranked genes by the density of FLC-specific enhancers that contain one or more FOSL2/JUN or CREB motifs. On the basis of these criteria, *SLC16A14* has the highest density of FLC-specific enhancers containing FOSL2/JUN motifs (Fig. 3D). *CA12* is also highly ranked in this version of the analysis. *TNRC6C* and *LINC00473* have the highest and second highest density of FLC-specific enhancers containing CREB motifs, respectively (Fig. 3E).

Next, we found that genes with at least one nearby FLC-specific enhancer containing either a FOSL2/JUN or CREB motif are over-represented in the MAPK signaling pathway (KEGG 2016, p=0.032, Fisher’s exact test) and significantly enriched for MAPK1 (ERK2) targets and interacting proteins (PPI Hub Proteins, p=5.4 × 10^−6^, Fisher’s exact test). We then ranked genes according to the density of FLC-specific enhancers containing both FOSL2/JUN and CREB motifs (Fig. 3F), and observed that genes with at least one nearby FLC-specific enhancer containing both FOSL2/JUN and CREB motifs are enriched for MAPK1 and MAPK3 (ERK1) targets and interacting proteins (Fig. 3G). Taken together, these results suggest that the regulation of genes such as *CA12, SLC16A14*, and *LINC00473* by FLC-specific enhancers may be mediated through FOSL2/JUN and CREB and that these transcription factors may contribute to and/or result from dysregulated MAPK signaling.

### Identification of FLC-specific enhancer target genes

Our gene window analyses (Fig. 3) identified genes that may be regulated by FLC-specific enhancers. However, nearby genes may have the same TREs within their windows. In order to more confidently link individual FLC-specific enhancers with putative gene targets, we correlated enhancer activity to gene transcription levels across all FLC tumors, as described previously (Corces et al., 2018). ChRO-seq allows us to quantify both enhancer activity and gene transcription from a single experimental dataset, thereby avoiding confounding variables that can arise when using multiple different assays. Correlations between transcriptional activity of enhancers and genes within 100 kb of each other were compared to a null distribution of inter-chromosomal enhancer-gene pairs to calculate p-values (Fig. 4A,B). Windows larger and smaller than 100 kb had reduced power to detect statistically significant gene-enhancer correlations (Fig. S3A). Using an FDR < 0.1, we linked 1697 FLC-specific enhancers to putative target genes (Fig. 4B,C). As expected, we observed that the frequency of predicted gene-enhancer links decreases with increasing distance between them (Fig. 4D). We found that most enhancers are linked to only one or two genes (mean = 1.33, Fig. 4E) and most genes are linked to only one or two enhancers (mean = 1.78, Fig. 4F). The top 5% of genes (in terms of enhancer connectivity) are each linked to at least 4 enhancers (Fig. 4G, Table S5). The top 5% includes *FAM19A5, LINC00473, VCAN, SLC16A14* (Fig. 4H) and *CA12* (Fig. 4I), which are putatively linked to 20, 9, 9, 9, and 6 FLC-specific enhancers, respectively (Fig. 4G-I; Fig. S4; Table S6). Interestingly, while *CA12* is not highly transcribed in NML, there is a substantial amount of ChRO-seq signal at the promoter (Fig. 4I), indicative of polymerase pausing. This is unlike what we observe at the *SLC16A14* locus (Fig. 4H), where there is no ChRO-seq signal in NML even at the TSS. This observation suggests that transcriptional pausing may be another mechanism that regulates *CA12* expression. Therefore, *CA12* may be poised for expression, whereas the *SLC16A14* locus is completely inactive in NML and dramatically rewired for activation in FLC.

**Figure 4.**
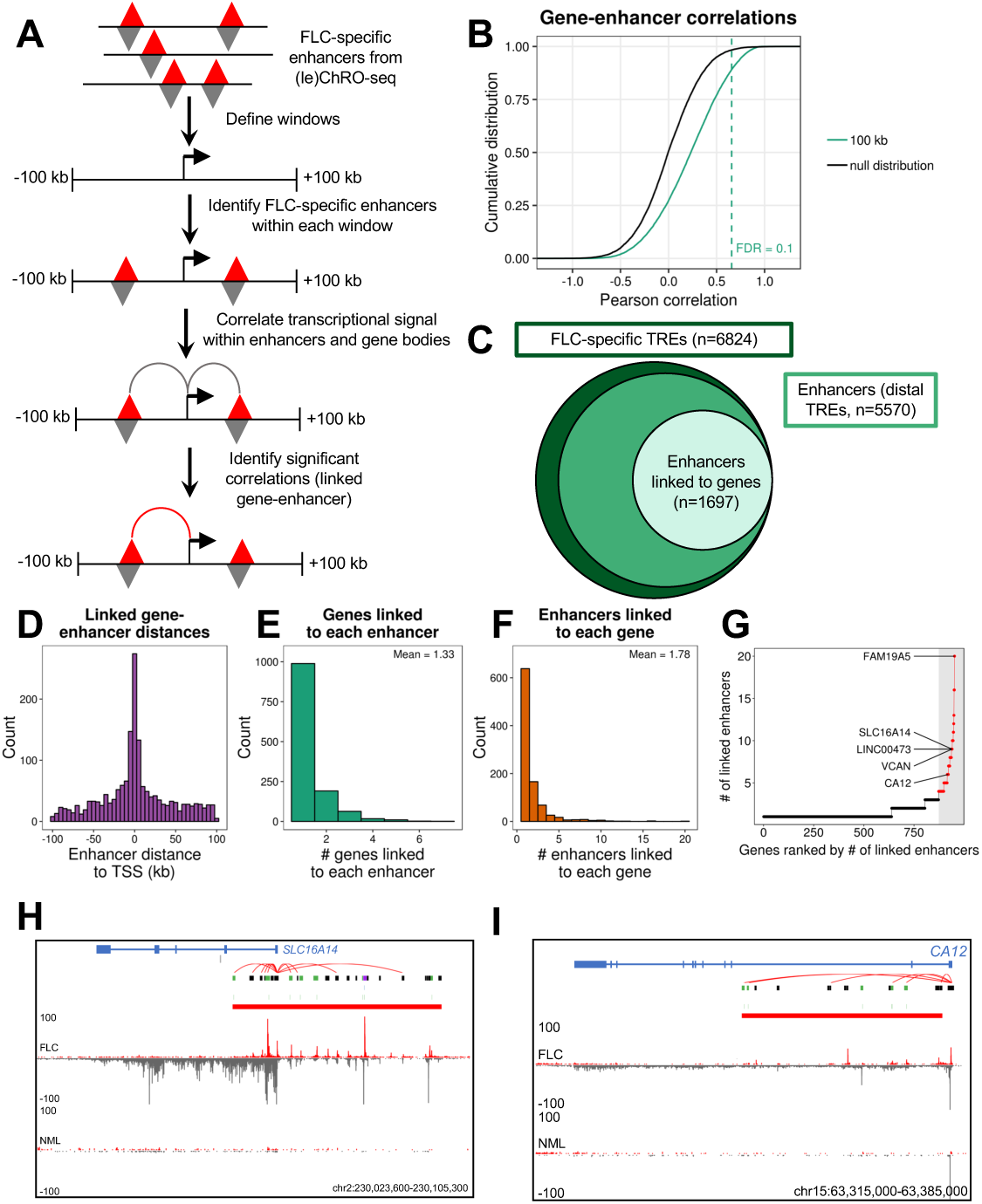
Identification of candidate target genes of individual enhancers. **(A)** Schematic of the approach used to link individual FLC-specific enhancers to candidate target genes. **(B)** Cumulative distribution function of the Pearson correlation coefficient from all gene-enhancer pairs within 100 kb and the null distribution (interchromosomal gene-enhancer pairs). **(C)** Plot showing the number of total FLC-specific TREs, enhancers, and enhancers linked to genes identified. **(D,E,F)** Histograms of the distance between linked genes and enhancers (D), number of gene targets linked to each enhancer (E), and number of enhancers linked to each target gene (F). **(G)** Ranked dot plot showing the number of enhancers linked to each target gene. Dots in red indicate the top 5% of genes based on number of linked enhancers (≥4 linked enhancers). **(H,I)** Genome snapshot of the *SLC16A14* (H) and *CA12* (I) loci. Computationally predicted gene-enhancer links, FLC-specific TREs, CREB motifs within FLC-specific TREs (blue), and FOSL2/JUN motifs within FLC-specific TREs (green) are shown below the gene diagram. FLC-specific TREs containing CREB motifs are shown in blue, those containing FOSL2/JUN motifs in green, those containing both CREB and FOSL2/JUN motifs in purple, and all others in black. Transcriptional signal from the plus and minus strand are shown in red and grey, respectively. FLC and NML show similar levels of paused polymerase for *CA12* (peak at TSS), but FLC has significantly more gene body transcription.

### Defining FLC-specific enhancer hotspots

Super enhancers are non-coding regions exhibiting unusually high transcriptional activity (Hnisz et al., 2013; Lovén et al., 2013; Pott and Lieb, 2015; Whyte et al., 2013) and are important regulators of cell identity (Hnisz et al., 2013; Whyte et al., 2013) and key cancer genes (Hnisz et al., 2013; Lovén et al., 2013). We used an algorithm analogous to ones previously used to define super enhancers from chromatin immunoprecipitation and sequencing (ChIP-seq) data (Lovén et al., 2013; Whyte et al., 2013) to identify clusters of enhancers that have remarkably high transcriptional activity in FLC, but not in NML (Fig. 5A, see Materials and Methods). Using FLC-specific enhancers as input, we identified 141 dense clusters with especially high FLC-specific transcriptional activity (Fig. 5B). Because these loci consist of only FLC-specific enhancers instead of all enhancers present in FLC, we refer to them as “FLC-specific enhancer hotspots” rather than FLC super enhancers.

**Fig. 5.**
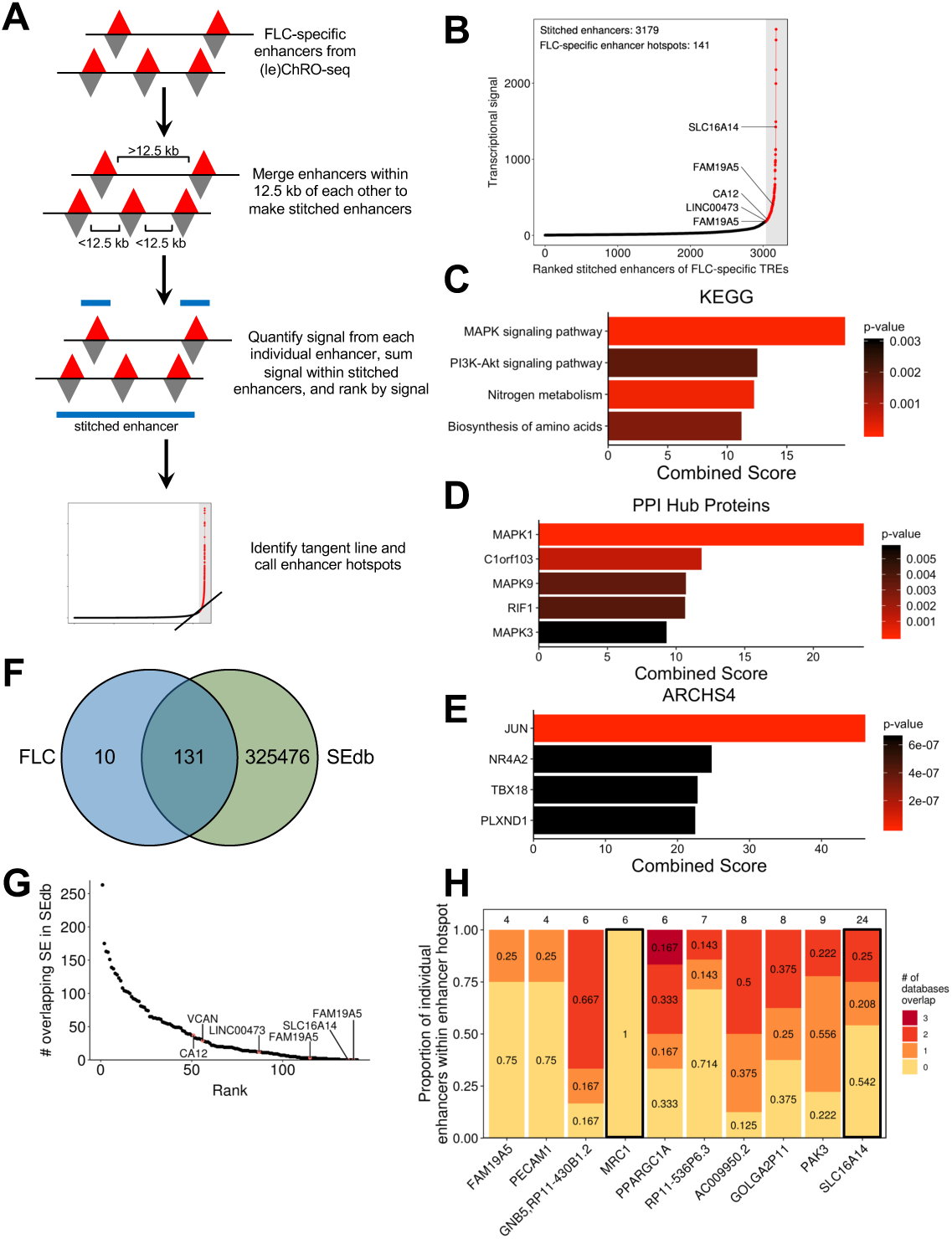
Identification of FLC-specific enhancer hotspots. **(A)** Schematic of the approach used to identify FLC-specific enhancer hotspots. **(B)** Total transcriptional signal from enhancers stitched from FLC-specific enhancers. Stitched enhancers are ranked based on transcriptional signal within FLC-specific enhancers. Points in red denote FLC-specific enhancer hotspots. **(C,D,E)** KEGG 2016 enrichment (C), PPI hub proteins (D), and ARCHS4 co-correlation (E) analysis of candidate target genes of FLC-specific enhancer hotspots. **(F)** Venn diagram showing the overlap between FLC-specific enhancer hotspots and all super enhancers in SEdb. The overlap displays the number of FLC-specific enhancer hotspots overlapping any super enhancer within SEdb rather the reciprocal comparison (n=5274). **(G)** Dot plot showing the number of overlapping super enhancers within SEdb for each FLC-specific enhancer. **(H)** Stacked bargraph examining the 10 FLC-specific enhancer hotspots that did not overlap with super enhancers in SEdb. Individual enhancers from each enhancer hotspot were examined for overlap with any enhancer from ENCODE, FANTOM5, and NIH Roadmap Epigenomics databases. The number above each bar indicates the total number of individual enhancers that comprise that enhancer hotspot. Bars outlined in black designate genes that have sufficient enhancer signal in individual unique enhancers to meet the threshold originally determined for FLC-specific enhancer hotspots.

As super enhancers have been shown to regulate key cancer drivers (Lovén et al., 2013), we linked each FLC-specific enhancer hotspot to the closest gene that is both robustly transcribed (ChRO-seq: TPM ≥ 25) and significantly increased in transcription (ChRO-seq: FDR < 0.05, log2fold change ≥ 1) in FLC compared to NML. Several of these genes have been consistently linked to FLC, including *CA12, SLC16A14, LINC00473, OAT, TMEM163*, and *TNRC6C*. Others, such as *FAM19A5*, have not previously been reported as dysregulated in FLC and represent novel candidate oncogenes. Enrichment analysis of genes linked to FLC-specific enhancer hotspots revealed significant over-representation in the MAPK signaling pathway (Fig. 5C) and enrichment for MAPK1 targets (Fig. 5D). Correlation analysis using the ARCHS4 database (Lachmann et al., 2018) demonstrated that these genes are also strongly linked to JUN expression (Fig. 5E). This finding is consistent with our previous analysis that JUN may be critical for the transcriptional regulation of these genes.

To determine the uniqueness of the FLC-specific enhancer hotspots, we compared them to previously identified super enhancers in SEdb (Jiang et al., 2019). This database contains 325,607 super enhancers from more than 540 human samples across over 240 cell and tissue types, including 14 from normal human liver, primary hepatocytes, and multiple cell lines including HepG2 and Huh7. We found that 10 FLC-specific enhancer hotspots are not present in any sample in SEdb. Notably, this includes those linked to *SLC16A14* and *FAM19A5* (Fig. 5F,G; Table S5), indicating that the mechanisms of transcriptional regulation of these genes may be unique in FLC. FLC-specific enhancer hotspots associated with *LINC00473* and *CA12* showed minimal intersection with super enhancers in SEdb (Fig. 5G), overlapping 12 and 37 super enhancers, respectively. Additionally, 72 FLC-specific enhancer hotspots, including those associated with *LINC00473* and *CA12*, did not overlap with super enhancers from any of the 14 liver-derived samples, which include healthy liver and hepatocellular carcinoma cell lines.

To further investigate the uniqueness of the 10 FLC-specific enhancer hotspots not present in SEdb, we cross-referenced these enhancer hotspots to individual enhancers identified by the ENCODE, FANTOM5, and NIH Roadmap Epigenomics consortia (see Materials and Methods). Only one FLC-specific enhancer hotspot (located near *MRC1*) did not overlap any individual enhancers recorded in the three databases mentioned above, suggesting this enhancer hotspot is truly unique to FLC (Fig. 5H). We also recalculated the signal for the same 10 FLC-specific super enhancers including only those individual enhancers that do not overlap with any enhancers from the three databases. Only the revised stitched enhancers close to *MRC1* and *SLC16A14* exhibited enough transcriptional activity (ChRO-seq signal) to still meet the threshold set in our original analysis for an enhancer hotspot. These results suggest that there is a substantial amount of enhancer activity that is potentially completely unique to FLC at the *MRC1* and *SLC16A14* loci.

### High-confidence candidate oncogenes in FLC

The genes significantly correlated with FLC-specific enhancers and those significantly associated with FLC-specific enhancer hotspots represent those genes that likely drive key oncogenic attributes of FLC cells. To further refine this list and identify high-confidence candidates, we integrated our ChRO-seq results with RNA-seq data from 23 FLC samples (Table S1, S6), ten of which also underwent ChRO-seq analysis. We first selected genes that were identified as both significantly correlated with FLC-specific enhancers (Fig. 4G) and linked to FLC-specific enhancer hotspots (Fig. 5B). These genes were then filtered to identify genes that are both highly transcribed (ChRO-seq: TPM ≥ 25, fold change ≥ 5, FDR < 0.2) and expressed (RNA-seq: normalized counts ≥ 100, fold change ≥ 5, FDR < 0.2) in FLC relative to NML. Integration of both ChRO-seq and RNA-seq data ensures that the transcriptional changes of these genes are maintained at the steady state RNA level. The final list harbored 16 genes (Fig. 6A, Table S6). We noticed that over half of these genes have been previously implicated in drug resistance (*CA12, COL4A1, HSPA1B, IRF4, KIF26B, LINC00473, SLC16A14, TESC*, and *VCAN*) and 10 of them are connected to elevated MAPK/ERK activity (*BACE2, CA12, COL4A1, FAM19A5, HSPA1B, IRF4, KIF26B, LINC00473, TESC*, and *VCAN*; Table S6).

**Fig. 6.**
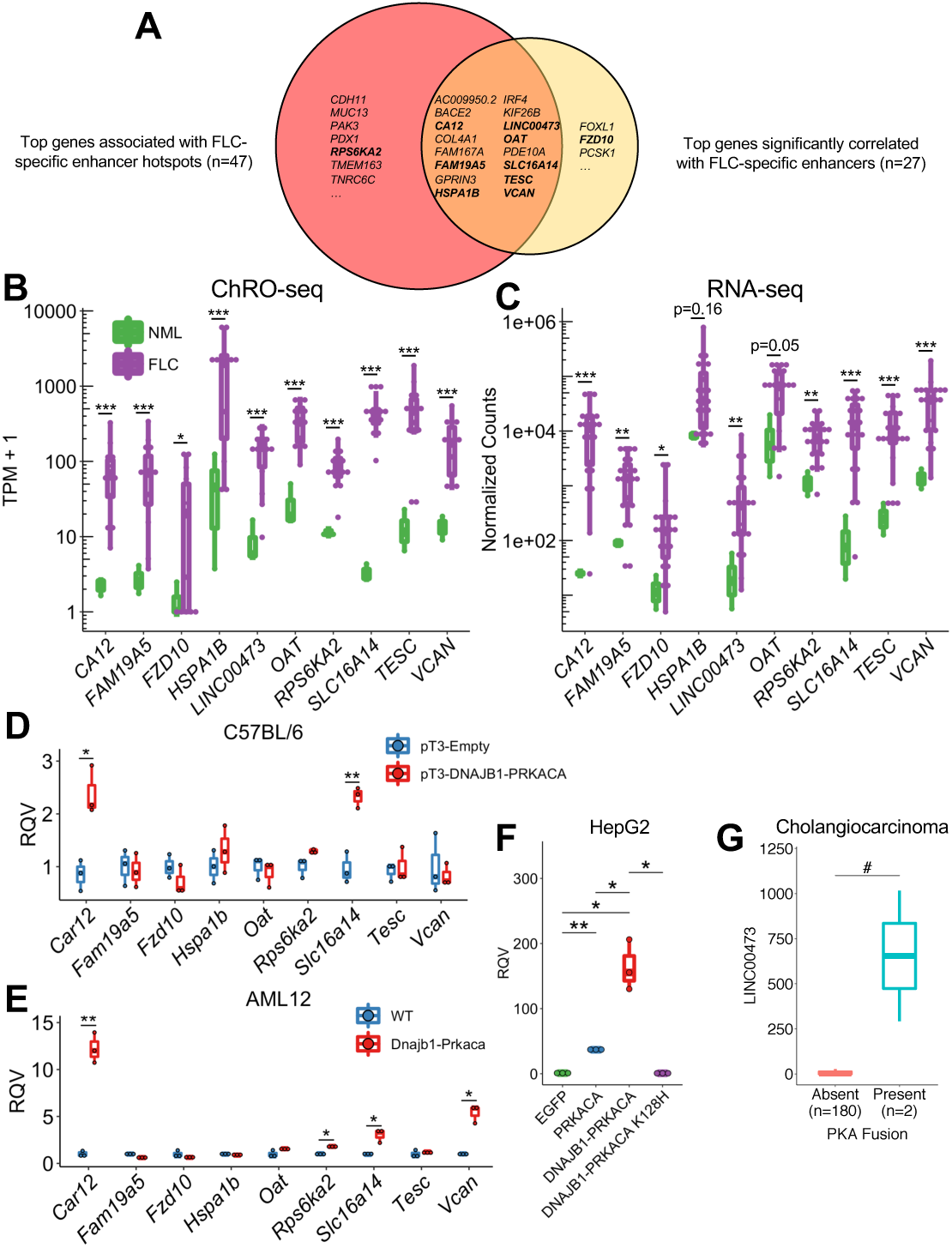
Candidate FLC oncogenes are transcriptionally dysregulated. **(A)** Venn diagram showing overlap of genes linked to FLC-specific enhancer hotspots and genes significantly correlated to FLC-specific enhancers. Genes in bold indicate those shown in panels B,C. **(B,C)** Boxplots showing transcription (B) and RNA expression (C) in FLC compared to NML. *p<0.05, **p<0.01, ***p<0.001 (Wald test, DESeq2). **(D)** RNA expression in liver tissue and tumors expressing empty (pT3-Empty) and fusion-containing (pT3-DNAJB1-PRKACA) transposon, respectively. **(E)** RNA expression in WT AML12 cells and a clone expressing DNAJB1-PRKACA. **(F)** RNA expression of *LINC00473* following expression of various *PRKACA* constructs. **(G)** RNA expression of *LINC00473* in cholangiocarcinoma samples with and without PKA fusions. *p<0.05, **p<0.01 (two-sided Welch’s t-test), ^#^p<0.05 (two-sided Mann-Whitney U test).

We selected eight of these genes (*CA12, FAM19A5, HSPA1B, LINC00473, OAT, SLC16A14, TESC*, and *VCAN*) for further investigation. We chose *SLC16A14* and *LINC00473* because they have high densities of FLC-specific enhancers with FOSL2/JUN (Fig. 3D) and CREB motifs (Fig. 3E), respectively; *FAM19A5* because it has the greatest number of significantly correlated enhancers (Fig. 4G); *TESC* and *VCAN* because they have previously been strongly implicated in drug resistance (Lee et al., 2018; Li et al., 2017; Man et al., 2014), a salient feature of FLC (Maniaci et al., 2009; Torbenson, 2012); *CA12* and *OAT* because we have previously identified them as candidate markers of FLC (Dinh et al., 2017) and because *CA12* is a prominent mediator of drug resistance in other cancer types (Boyd et al., 2017; Doyen et al., 2013; Kopecka et al., 2016; Yoo et al., 2010); and *HSPA1B* because it encodes a member of the heat shock protein 70 (HSP70) family, which has been shown to interact with the DNAJB1-PRKACA fusion found in FLC (Turnham et al., 2019). We also selected one gene that only appears in the FLC-specific enhancer hotspot analysis (*RPS6KA2*) and another from the gene-enhancer correlation (*FZD10*) analysis for further investigation (Fig. 6A).

We compared the expression of these genes in FLC to other cancer types within The Cancer Genome Atlas (TCGA) and found that *SLC16A14* is more highly expressed in FLC than any other cancer type (Fig. S5). *FAM19A5, LINC00473, CA12, TESC*, and *VCAN* are also highly expressed in FLC, in addition to several other cancer types within TCGA (Fig. S5). We also compared *SLC16A14* expression in FLC compared to normal tissues within the Genotype-Tissue Expression (GTEx) database and found substantially higher levels of *SLC16A14* in FLC than any other normal tissue (Fig. S5H).

### *SLC16A14, CA12, LINC00473, RPS6KA2*, and *VCAN* are responsive to DNAJB1-PRKACA

While the genes we selected are over-transcribed (Fig. 6B) and over-expressed (Fig. 6C) in FLC relative to NML, it is unclear whether this dysregulation is directly caused by the DNAJB1-PRKACA fusion. To determine whether the DNAJB1-PRKACA fusion is sufficient to perturb these genes of interest, we took advantage of two murine models of FLC. In the first model, a transposon expressing human *DNAJB1-PRKACA* is introduced into the livers of C57BL/6 mice by hydrodynamic tail vein injection, forming FLC-like liver tumors (Kastenhuber et al., 2017). We examined the expression of our genes of interest in the resulting liver tumors compared to livers from mice injected with an empty vector control. *Car12* (the mouse homolog of human *CA12*) and *Slc16a14* displayed significantly higher expression of samples expressing *DNAJB1*-*PRKACA* compared to control (Fig. 6D). The second model is the AML12 cell line that has undergone CRISPR/Cas9 gene editing, resulting in a heterozygous deletion analogous to the endogenous event in humans, leading to the formation of the *Dnajb1*-*Prkaca* fusion (Dinh et al., 2019; Turnham et al., 2019). When we examined the expression of the genes of interest in AML12 cells expressing *Dnajb1*-*Prkaca*, we found that *Car12* and *Slc16a14*, as well as *Rps6ka2* and *Vcan*, are significantly elevated compared to wild-type (WT) controls (Fig. 6E).

As *LINC00473* is a primate-specific lncRNA (Reitmair et al., 2012), we used an alternative non-murine model to study its regulation. Specifically, we stably over-expressed the fusion in the HepG2 human hepatoma cell line using a lentiviral system. *DNAJB1-PRKACA* expression dramatically increased *LINC00473* expression compared to an enhanced green fluorescent protein (*EGFP*) control (Fig. 6F). *WT PRKACA* also increased *LINC00473* expression. However, the magnitude of *LINC00473* induction was significantly larger with the fusion compared to *WT PRKACA*, indicating that something other than canonical PKA activity (possibly the DNAJB1 domain) is relevant for robust induction of LINC00473. Importantly, stable expression of a kinase-dead mutant of *DNAJB1-PRKACA* (K128H) did not increase *LINC00473* expression (Fig. 6F), indicating that the kinase activity of the fusion is necessary for induction of expression. To determine if *LINC00473* might be responsive to PKA fusions in other contexts, we examined a cholangiocarcinoma dataset (Nakamura et al., 2015) that characterized tumors with fusions involving *PRKACA* or *PRKACB* with ATPase Na+/K+ transporting subunit beta 1 (*ATP1B1*). Interestingly, the exons of *PRKACA* retained in the *ATP1B1-PRKACA* fusion are the same as in *DNAJB1-PRKACA*. Using RNA-seq data generated for this dataset, we examined the relationship between PKA fusions and the expression of *LINC00473*. Tumors with PKA fusions demonstrated significantly higher expression of *LINC00473* than tumors without PKA fusions (Fig. 6G), indicating that *LINC00473* is responsive to PKA activity in alternative fusion events. Our results suggest that DNAJB1-PRKACA is sufficient to perturb the expression of *CA12, SLC16A14, VCAN, RPS6KA2*, and *LINC00473* in the specific disease models we used and that this regulation is dependent upon (at least for *LINC00473*) the kinase activity of DNAJB1-PRKACA.

### Suppression of SLC16A14 or CA12, either alone or in combination with a MAPK inhibitor, reduces viability of FLC cell models

Our results thus far suggest that MAPK signaling regulates FLC pathogenesis (Fig. 3G, 5C,D). To confirm that MAPK signaling is overactive in FLC, we measured the levels of phosphorylated MEK and ERK in WT and *Dnajb1-Prkaca*-expressing AML12 cells (Fig. 7A,B). As expected, we observed dramatically increased phospho-MEK and phospho-ERK in cells expressing the fusion compared to WT cells. Treatment of AML12 cells expressing *Dnajb1-Prkaca* with the MEK inhibitor cobimetinib resulted in a dose-responsive decrease in cell viability (Fig. 7C-F). Both SLC16A14 and CA12 (*Car12*), which we identified as prominent FLC-enhancer-hotspot associated genes, have been implicated in drug resistance in other cancers (Doyen et al., 2013; Januchowski et al., 2014; Kopecka et al., 2016), and CA12 has been reported previously as a mediator of the effects of the MAPK pathway (Hsieh et al., 2010). Knockdown of *Slc16a14* by siRNA dramatically reduced viability of AML12 cells expressing *Dnajb1-Prkaca*, and also increased the potency of cobimetinib (Fig. 7C,D). Knockdown of *Car12* did not have much of an effect on its own, but in combination with cobimetinib did significantly reduce cell viability compared to cobimetinib alone (Fig. 7E,F, Fig. S6A).

**Fig. 7.**
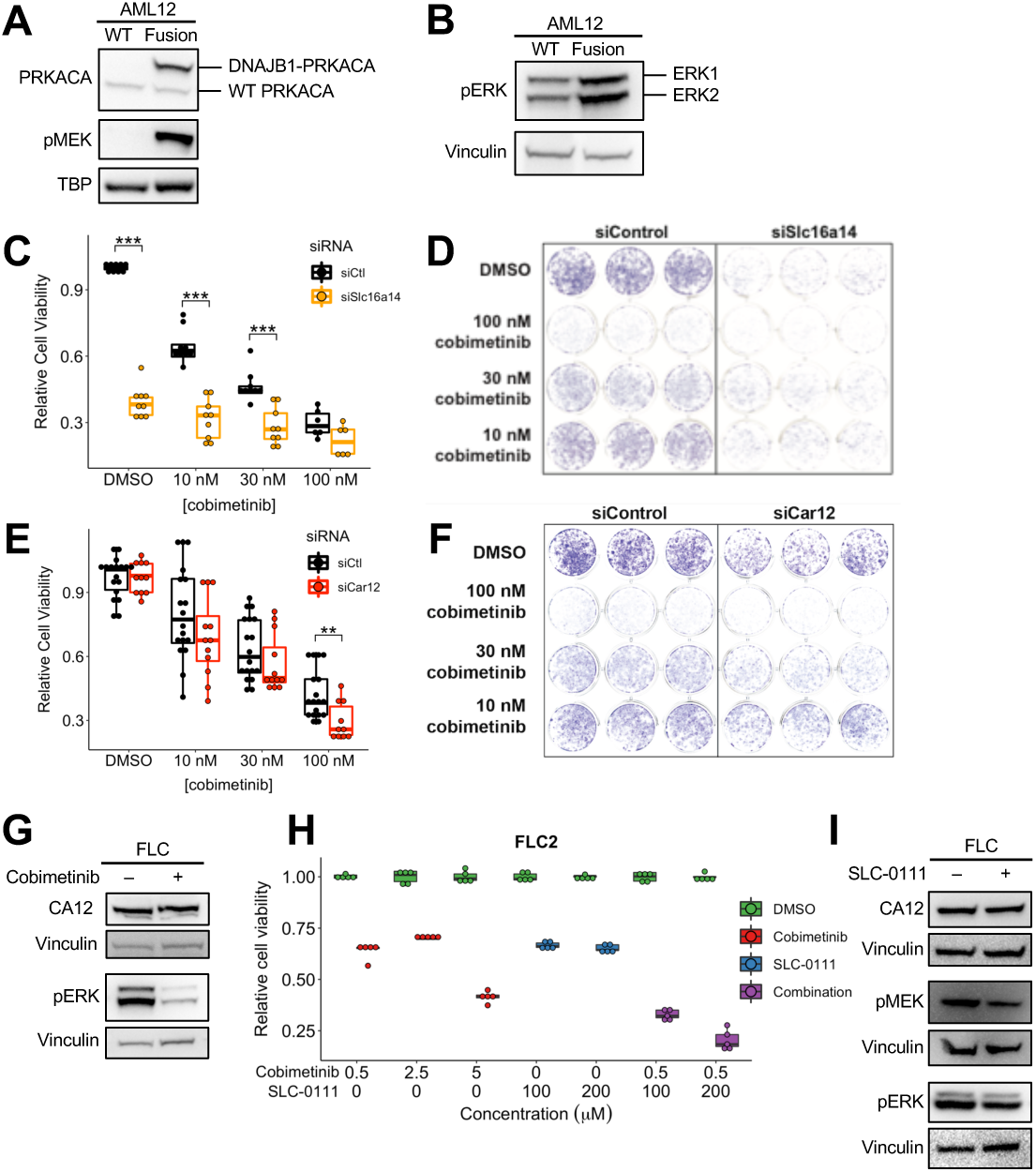
The MAPK and SRC signaling pathways are dysregulated in FLC. **(A,B)** Western blot in WT AML12 cells and AML12 cells expressing the DNAJB1-PRKACA fusion demonstrating elevated MEK (A) and ERK (B) phosphorylation in cells expressing the fusion. **(C,E)** Cell viability quantified by crystal violet staining in AML12 cells expressing the DNAJB1-PRKACA fusion. Cells were treated with a siRNA targeting *Slc16a14* (C), *Car12* (E), or a control siRNA and multiple concentrations of the MEK inhibitor cobimetinib. **(D,F)** Representative wells for cells treated with multiple concentrations of cobimetinib and siRNA targeting *Slc16a14* (D), *Car12* (F), or a control siRNA and stained with crystal violet. **(G)** Western blot in FLC cells treated with 2.5 μM cobimetinib and probed with antibodies detecting CA12 or phosphorylated ERK. **(H)** Cell viability quantified by CellTiter-Glo in FLC cells treated with cobimetinib alone or the combination of cobimetinib and SLC-0111. All comparisons between DMSO and treatment were statistically significant (p<0.01). **(J)** Western blot in FLC cells treated with 200 μM SLC-0111 and probed with antibodies detecting CA12, phosphorylated MEK, or phosphorylated ERK. **p<0.01, ***p<0.001 (two-sided Mann-Whitney U test).

We next tested whether inhibition of MAPK signaling reduces viability of human FLC tumor cells. First, we derived a primary human FLC cell line from a patient-derived xenograft (PDX) model by optimizing previously described protocols (Dinh et al., 2019; Liu et al., 2017, see Materials and Methods). Treatment with cobimetinib reduced ERK phosphorylation as expected (Fig. 7G), and significantly decreased FLC cell viability (Fig. 7H), but only at higher doses of the drug. This effect was confirmed across three distinct derivations of the FLC cell line (Fig. 7H, Fig. S6B), each of which was positive for *DNAJB1-PRKACA* expression. Our results suggest that FLC cells are susceptible to MAPK inhibition, however the doses needed for a cytotoxic effect indicate an intrinsic level of drug resistance in these cells.

We hypothesized that inhibition of CA12 in FLC cells may mitigate the intrinsic drug resistance because of the results in the AML12 cells expressing *Dnajb1-Prkaca* (Fig. 7E) and because an effective CA12 inhibitor, SLC-0111, is currently in clinical trials as combination therapy for metastatic pancreatic cancer. We found that pharmacological suppression of CA12 with SLC-0111 in FLC cells significantly enhances the potency of cobimetinib at the 500 nM dose (Fig. 7H, Fig. S6B). Again, this effect was confirmed across three distinct derivations of the FLC cell line. We quantified the interaction between cobimetinib and SLC-0111 using the combination index (CI). SLC-0111 and cobimetinib combination treatment resulted in a synergistic response (CI = 0.00686 (0.5 μM cobimetinib + 100 μM SLC-0111) and 0.000791 (0.5 μM cobimetinib + 200 μM SLC-0111) in FLC2, where CI < 1 indicates synergy). To determine whether CA12 interacts with the MAPK signaling pathway, we inhibited either MEK or CA12. Cobimetinib treatment resulted in decreased ERK phosphorylation as expected, but had no effect on CA12 expression (Fig. 7G). However, treatment with SLC-0111 reduced phosphorylated MEK and ERK levels (Fig. 7J), suggesting that CA12 may function, at least in part, upstream of the MAPK signaling pathway. Taken together, our results suggest that the MAPK signaling pathway is dysregulated in FLC and inhibition of this pathway in combination with pharmacological suppression of CA12 or inhibition of SLC16A14 represent exciting candidate molecular therapeutic approaches.

## Discussion

Fibrolamellar carcinoma is a devastating cancer affecting young adults with limited treatment options. Thus, there remains a great need to identify potential therapeutic targets. Here we have analyzed primary FLC tumors and matched NML samples to map the unique enhancer landscape of FLC. One of the goals of this study was to discover master regulators of dysregulated gene expression and signaling in FLC. We took advantage of the new technique (le)ChRO-seq (Chu et al., 2018), which allowed us to perform run-on sequencing on frozen primary tumors. A notable advantage of (le)ChRO-seq is that it allows quantification of both enhancer activity and gene transcription from a single experiment, thereby avoiding possible confounders associated with using multiple different assays.

Our finding that FLC-specific TREs are enriched in motifs of FOSL2/JUN and CREB is noteworthy for several reasons. First, CREB is a well-validated substrate of wild-type PKA (Shaywitz and Greenberg, 1999) and has previously been shown to be hyperphosphorylated in FLC compared to adjacent liver (Xu et al., 2014). Second, CREB transcriptionally activates LINC00473 (Chen et al., 2016, 2018; Reitmair et al., 2012) and represses miR-375 (Keller et al., 2012), the latter of which is a candidate tumor suppressor in FLC (Dinh et al., 2019). Third, CREB and AP-1, a heterodimer consisting of FOS and JUN subunits, have been shown to regulate each other (Ma et al., 2014; Sanyal et al., 2002). Finally, CREB and AP-1 are both activated by the MAPK signaling pathway (Ginty et al., 1994; Karin, 1995; Wu et al., 2001; Xing et al., 1996), which is dysregulated in FLC (Turnham et al., 2019).

Our analysis reveals that transcription at distal TREs (i.e. enhancers) stratify FLC from NML samples better than transcription at proximal TREs or gene bodies, similar to previous reports (Corces et al., 2016, 2018). These findings are consistent with other studies (Franco et al., 2018; Van Groningen et al., 2017; Ooi et al., 2016) that have shown enhancer activity is more sensitive than gene expression for sample classification (e.g., tumor-normal, tumor subtypes, cell types). Unlike other methods to identify enhancers, such as assay for transposase-accessible chromatin using sequencing (ATAC-seq) or chromatin immunoprecipitation and sequencing (ChIP-seq) for enhancer-associated histone modifications or proteins, ChRO-seq identifies active regulatory elements as well as actively transcribed genes. This advantage allowed us to correlate enhancer and gene transcription within the same assay, as well as to provide a more direct output of enhancer function (transcription) than steady state RNA levels, which reflect cumulative effects of transcriptional, co-transcriptional, and post-transcriptional regulatory processes.

To identify putative targets of FLC-specific enhancers, we employed computational approaches based on enhancer density and signal correlation. We identified 141 FLC-specific enhancer hotspots, defined as dense clusters of enhancers with especially high transcriptional activity (similar to the concept of super enhancers). Comparing these loci with previously characterized super enhancers in SEdb, we found that 10 FLC-specific enhancer hotspots are completely unique and do not overlap with any known super enhancer. Furthermore, 72 of the enhancer hotspots do not overlap with super enhancers from any previously assayed liver tissue or liver-derived cells. As super enhancers have been shown to regulate cell identity and oncogenes (Hnisz et al., 2013; Lovén et al., 2013), our findings suggest that these FLC-specific enhancer hotspots may regulate genes important in the formation, progression, and/or maintenance of this cancer. Super enhancers are regulated by the bromodomain protein BRD4 (Lovén et al., 2013), and BRD4 inhibitors such as JQ1 (Filippakopoulos et al., 2010) have been shown to disrupt super enhancer function (Gryder et al., 2017; Lovén et al., 2013; Mack et al., 2018; Peeters et al., 2015). Pharmacological disruption of the FLC-specific enhancer hotspots we have described here may represent an alternative therapeutic approach for FLC.

We have computationally identified gene-enhancer interactions in this study. However, future experiments will be important to confirm these interactions experimentally and carefully dissect the 3-D regulatory interactions in FLC. For example, methods based on chromosome conformation capture, such as Hi-C, can identify and confirm global chromosomal interactions, while methods based on luciferase assays, such as self-transcribing active regulatory region sequencing (STARR-seq) or other massively parallel reporter assays, can validate the regulatory activity of identified enhancers. More refined dissection of individual or subsets of enhancers using CRISPR/Cas9 genome editing will confirm functional gene-enhancer links and identify the specific nucleotides within enhancers that are essential for such regulation. Furthermore, our ChRO-seq experiments were conducted with primary tumors, which contain a heterogenous mixture of cells. Our results therefore represent the aggregate results across all cell types present. Going forward, single cell experiments will be necessary to dissect the role of transcriptional regulatory networks in tumor, immune, parenchymal, and other cell types within primary tumors.

Integrative analyses of ChRO-seq and RNA-seq data identified 16 high-confidence candidate oncogenes in FLC. Some of these genes, including *LINC00473* and *CA12* have been characterized in other cancers previously. *LINC00473* is known to be overexpressed in FLC (Dinh et al., 2017) and other cancers (Chen et al., 2016, 2018; Shi et al., 2017), and promotes chemotherapeutic resistance in colon cancer (Wang et al., 2018) and head and neck squamous cell carcinoma (Han et al., 2018). *CA12* has been linked to drug resistance in multiple cancer types (Boyd et al., 2017; Doyen et al., 2013; Kopecka et al., 2016) and we have previously demonstrated that it is overexpressed in FLC (Dinh et al., 2017). Other genes, including *SLC16A14* and *FAM19A5*, are relatively understudied and merit deeper investigation.

Harnessing well-characterized inhibitors of some of these genes may represent a potentially expedient avenue for new FLC therapeutics. One notable example is an inhibitor of CA12, SLC-0111, currently in clinical trials for metastatic pancreatic cancer (see below). Most of the genes we identified here are not currently targeted by drugs or inhibitors; nonetheless, alternative strategies exist to target cells uniquely expressing these genes. Drugs conjugated to antibodies or aptamers that can bind to cell surface proteins that are specific to cells of interest, such as SLC16A14 in FLC cells, represent an emerging strategy for targeting tumor cells. Examination of *SLC16A14* expression in the Genotype-Tissue Expression (GTEx) database demonstrated substantially higher expression in FLC tumors than any other normal tissue, indicating that it may be a useful molecular beacon for such approaches. For non-cell surface proteins, this method can be modified by engineering T-cells that recognize the major histocompatibility complex (MHC) class I presenting specific peptides from the protein of interest. Leveraging the knowledge gained in this study to develop new targeted therapeutic approaches remains an important goal.

Our analyses show that a large proportion of the genes that are associated with FLC-specific enhancers function in drug resistance, including *LINC00473* (Han et al., 2018; Wang et al., 2018), *VCAN* (Li et al., 2017), and *CA12* (Boyd et al., 2017; Doyen et al., 2013; Kopecka et al., 2016; Yoo et al., 2010). This suggests that high expression of these genes may be responsible for the strong drug resistance phenotype observed in FLC. Thus, a viable therapeutic strategy to combat drug resistance may be to combine inhibitors of drug resistance genes with presently available therapeutics. For example, ongoing clinical trials for metastatic pancreatic cancer are focused on combining SLC-0111, a CA12 inhibitor, with the standard therapeutic gemcitabine (ClinicalTrials.gov Identifier: NCT03450018).

In addition, we have demonstrated that *CA12, SLC16A14, VCAN*, and *RPSK6A2* are responsive to DNAJB1-PRKACA in at least one of two different genetically engineered murine models of FLC. However, we did not notice a significant induction of expression for several of the other FLC enhancer-hotspot associated genes that we tested, even though they are over-expressed in primary FLC tumors. There are several possible explanations for this observation. First, the murine models we used might have species-specific differences in gene regulation compared to human. Second, these genes might not be directly downstream of DNAJB1-PRKACA, but induced due to another process during tumor initiation or progression. For example, certain genes may be induced during or in response to the development of tumor fibrosis within FLC tumors. Although this is a distinctive feature of primary human FLCs, current mouse models lack this characteristic, providing a possible explanation for the observed differences. While this is indicative of a critical need for better model systems, our results using two murine models suggest that at least *CA12, SLC16A14, VCAN*, and *RPS6KA2* are responsive to DNAJB1-PRKACA. Alternative models may be necessary to study the remaining genes. For example, since *LINC00473* is a primate-specific lncRNA, we used the HepG2 cell line stably expressing *DNAJB1-PRKACA* to demonstrate *LINC00473* is responsive to the fusion.

Finally, we demonstrated that the MAPK signaling pathway is dysregulated in FLC. While the cell line we have derived and used here is, to our knowledge, the only published FLC cell line, it is not without its limitations. As the FLC cells are derived from a PDX model and can be co-cultured with irradiated mouse fibroblasts, we observe a murine component in each derived cell line (see Materials and Methods). However, experiments across multiple cell line derivations with varying degrees of murine component produced similar results (Fig. 7H,I, Fig. S6B,C). These observations underscore the need for additional models of FLC, including new cell lines. The cell lines described here are all derived from the only published FLC PDX model (Oikawa et al., 2015), therefore derivations from additional PDX models will be important as they become available. In each cell line derivation, inhibition of the MAPK pathway significantly reduced cell viability, but most effectively in the micromolar range, suggesting these cells exhibit intrinsic drug resistance similar to what has been reported in patients. Combination treatment of MEK inhibitor with the CA12 inhibitor SLC-0111 resulted in enhanced potency compared to MEK inhibitor alone indicating that CA12 inhibition in combination with additional therapeutics might be a viable treatment strategy for FLC. Importantly, inhibitors targeting multiple members of the MAPK cascade are in clinical use for the treatment of other cancers and SLC-0111 is currently in clinical trials for metastatic pancreatic cancer. Repurposing these therapeutics for FLC patients may provide more effective treatments than the limited options currently available. Although we have demonstrated that inhibition of CA12 enhances the potency of MEK inhibitors, inhibition of other FLC enhancer hotspot-associated genes that are potentially involved in drug resistance, such as *SLC16A14, TESC*, or *VCAN*, may provide additional therapeutic benefit. Indeed, knockdown of *Slc16a14* in AML12 cells expressing *Dnajb1-Prkaca* both enhances the potency of cobimetinib and demonstrates substantial cytotoxic activity alone.

In sum, we have used ChRO-seq to map the transcriptional and enhancer landscape of FLC. The genome-scale information provided by ChRO-seq allowed us to identify candidate master transcriptional regulators of FLC and novel candidate FLC oncogenes, demonstrating the power of such genomic approaches. Follow-up functional studies in AML12 cells expressing *Dnajb1-Prkaca* and a newly derived human FLC cell culture revealed new candidate therapeutic strategies for FLC.

## Materials and Methods

### Lead Contact and Materials Availability

Further information and requests for resources and reagents should be directed to and will be fulfilled by the Lead Contact, Praveen Sethupathy.

### Experimental Model and Subject Details

#### Human liver samples

Informed consent was obtained from all human subjects. Samples were collected according to Institutional Review Board protocols 1802007780, 1811008421 (Cornell University) and/or 33970/1 (Fibrolamellar Cancer Foundation) and provided by the Fibrolamellar Cancer Foundation. Importantly, some samples come from the same patient (Table S2).

#### Animals

Samples from C57BL/6N mice were obtained for a previous study (Dinh et al., 2019) and remaining samples were used for this study. Briefly, female 6-10 week old C57BL/6N mice were subjected to hydrodynamic tail-vein injection with sterile 0.9% NaCl and 20 μg transposon and CMB-SB13 transposase (1:5 molar ratio). The transposon plasmid expressed human DNAJB1-PRKACA or an empty control. 3,5-diethoxycarbonyl-1,4-dihydrocollidine (DDC, 0.1%) diet was administered after tail-vein injection.

#### Cell lines

HepG2 expressing DNAJB1-PRKACA and EGFP have been previously described (Dinh et al., 2019). HepG2 cells were grown in Dulbecco’s Modified Eagle Media (DMEM) containing 1 g/L glucose (Thermo Fisher Scientific) supplemented with 10% fetal bovine serum (Thermo Fisher Scientific), 1% GlutaMAX (Thermo Fisher Scientific), 110 mg/L sodium pyruvate, and 1% penicillin-streptomycin (Thermo Fisher Scientific). The *DNAJB1-PRKACA* K128H kinase-dead mutant was cloned using the QuikChange II XL Site-Directed Mutagenesis Kit (Agilent Technologies) using the following primers (5’-CCTTCTGTTTGTCGAGGATATGCATGGCATAGTGGTTCCCG-3’ and 5’-CGGGAACCACTATGCCATGCATATCCTCGACAAACAGAAGG-3’). PCR products were cloned into the pCR-Blunt II-TOPO vector (Thermo Fisher Scientific) and subcloned into the pLV-EF1a-IRES-Puro vector (Addgene plasmid #85132, gift from Tobias Meyer). For lentivirus production, HEK293/T17 cells were transfected with DNAJB1-PRKACA K128H plasmid along with psPAX2 (Addgene plasmid #12260, gift from Didier Trono) and pMD2.G (Addgene plasmid #12259, gift from Didier Trono) to produce lentiviral particles, which were concentrated using Lentiviral-X Concentration (Takara Bio USA, Mountain View, CA) per manufacturer’s protocol. HepG2 cells were transduced with varying concentrations of lentivirus for 24 hours and selected with 2 μg/mL puromycin (Thermo Fisher Scientific) for 4 days. Selectable cells treated with the lowest concentration of virus were used for further passaging and experiments to obtain a majority of cells with 1 viral integration.

WT AML12 cells and AML12 cells expressing the DNAJB1-PRKACA fusion have been previously described (Dinh et al., 2019; Turnham et al., 2019). They were cultured in DMEM/F12 supplemented with 10% FBS, 0.04 μg/mL dexamethasone, 0.1% gentamicin, 1 μg/mL recombinant human insulin, 0.55 μg/mL human transferrin, and 0.5 ng/mL sodium selenite.

FLC cells were previously described (Dinh et al., 2019). Lines 3 and 18 were grown in complete F medium according to a previously published protocol (Liu et al., 2017). Line 2 was grown similarly to lines 3 and 18 with three minor modifications. First, F media was conditioned by irradiated mouse embryonic fibroblasts for 3 days prior to use. FLC cells were cultured in conditioned F media without irradiated fibroblasts. Second, R-spondin conditioned media was added to complete F media to 10% volume. R-spondin conditioned media was produced by culturing HEK293T cells expressing murine Rspo1 (gift from Alexander Nikitin lab) in conditioning media (DMEM supplemented with 1% GlutaMAX, 1% HEPES, and 1% penicillin-streptomycin) for 10 days. The supernatant was collected and filtered through a 0.22 μm filter prior to use. Third, the ROCK inhibitor Y-27632 was used a final concentration of 20 μM, increased from the original concentration of 10 μM (Liu et al., 2017). We detected the presence of murine cells in all three derived cell lines by RT-qPCR. All cell lines were cultured in 5% CO_2_ at 37°C.

## Method Details

### Chromatin run-on sequencing

ChRO-seq was performed as previously described (Chu et al., 2018; Mahat et al., 2016) with minor modifications. Length extension ChRO-seq (leChRO-seq) was performed identically to ChRO-seq except where indicated. Chromatin was isolated from pulverized frozen tissue in 1 mL 1X NUN buffer (20 mM HEPES, 7.5 mM MgCl_2_, 0.2 mM EDTA, 0.3 M NaCl, 1M urea, 1% NP-40, 1 mM DTT, 50 units/mL SUPERase In RNase Inhibitor (Thermo Fisher Scientific, Waltham, MA, AM2694), 1X Protease Inhibitor Cocktail (Roche, 11873580001)). For leChRO-seq, 50 units/mL RNase Cocktail Enzyme Mix (Thermo Fisher Scientific, AM2286) was substituted for SUPERase In RNase Inhibitor. Samples were vortexed for 1 minute, an additional 500 μL of 1x NUN buffer was added to each sample, and the samples were vortexed for an additional minute. Samples were incubated in an Eppendorf Thermomixer (Eppendorf, Hamburg, Germany) at 12°C and shaking at 2000 rpm for 30 minutes before centrifugation at 12,500 x g for 30 minutes at 4°C. Each sample was washed with 1 mL 50 mM Tris-HCl (pH 7.5) supplemented with 40 units/mL SUPERase In RNase Inhibitor (80 units/mL for leChRO-seq) and centrifuged at 10,000 x g for 5 minutes at 4°C. This wash step was repeated twice and samples were stored in 50 μL of chromatin storage buffer (50 mM Tris-HCl pH 8.0, 25% glycerol, 5 mM magnesium acetate, 0.1 mM EDTA, 5 mM DTT, and 40 units/mL SUPERase In RNase Inhibitor). Samples were loaded into a Bioruptor (Diagenode, Denville, NJ) and sonicated on the high power setting for a cycle time of 10 minutes, consisting of 10 cycles of 30 seconds on and 30 seconds off. Sonication was repeated as necessary to solubilize the chromatin and samples were stored at -80°C.

Following chromatin isolation, 50 μL of chromatin was mixed with 50 uL 2X run-on reaction mix (10 mM Tris-HCl pH 8.0, 5 mM MgCl_2_, 1 mM DTT, 300 mM KCl, 400 μM ATP, 0.8 μM CTP, 400 μM GTP, 400 μM UTP, 40 μM Biotin-11-CTP (Perkin Elmer, Waltham, MA, NEL542001EA), 100 ng yeast tRNA (VWR, Radnor, PA, 80054-306), 0.8 units/μL SUPERase In Rnase Inhibitor, 1% sarkosyl). The run-on reaction was performed at 37°C for 5 minutes at 700 rpm and stopped by adding 500 μL Trizol LS (Thermo Fisher Scientific, 10296-010) to the reaction. RNA samples were precipitated and resuspended in diethylpyrocarbonate (DEPC) treated water, heat treated at 65°C for 40 seconds, and digested on ice with 0.2N NaOH for 4 minutes. Base hydrolysis by NaOH was excluded from leChRO-seq protocols. Nascent RNA was purified with streptavidin beads (New England Biolabs (NEB), Ipswich, MA, S1421S) as previously described (Chu et al., 2018; Mahat et al., 2016). RNA was purified from beads using Trizol (Thermo Fisher Scientific, 15596-026) and 3’ adaptor ligation was performed with T4 RNA Ligase 1 (NEB, M0204S). Streptavidin bead binding was performed again following by 5’ decapping with RNA 5’ pyrophosphohydrolase (RppH, NEB M0356S). The 5’ end of the RNA molecule was phosphorylated with T4 polynucleotide kinase (PNK, NEB M0201S) and 5’ adaptor ligation was performed with T4 RNA Ligase 1. The 5’ adaptor contained a 6-nucleotide unique molecular identifier (UMI) to allow for bioinformatic detection and elimination of PCR duplicates. Streptavidin bead binding was performed again followed by reverse transcription using SuperScript IV Reverse Transcriptase (Thermo Fisher Scientific, 18090010). cDNA was amplified by PCR using the Q5 High-Fidelity DNA Polymerase (NEB, M0491S) to generate (le)ChRO-seq libraries. Libraries were sequenced (5’ single end) at the Biotechnology Research Center at Cornell University on the NextSeq500 (Illumina, San Diego, CA). Primer sequences used for (le)ChRO-seq library preparation are provided in Table S8.

### RNA-sequencing

Total RNA was isolated using the Total RNA Purification Kit (Norgen Biotek) per manufacturer’s instructions. RNA purity was quantified with the Nanodrop 2000 (Thermo Fisher Scientific, Waltham, MA) or Nanodrop One and RNA integrity was quantified with the Agilent 4200 Tapestation (Agilent Technologies, Santa Clara, CA) or Agilent BioAnalyzer. Libraries were prepared using the TruSeq Stranded mRNA Library Prep Kit (Illumina), the KAPA Stranded mRNA-Seq Kit (KAPA Biosystems, Wilmington, MA), or the NEBNext Ultra II Directional Library Prep Kit (New England Biolabs, Ipswich, MA). Sequencing was performed at the Biotechnology Research Center at Cornell University on the NextSeq500 (Illumina) or at the High-Throughput Sequencing Facility at the University of North Carolina at Chapel Hill on the HiSeq2500 (Illumina).

### Small RNA-sequencing

Small RNA sequencing was performed as previously described (Dinh et al., 2019). Briefly, reads were trimmed using Cutadapt and mapped to the genome using Bowtie (Langmead et al., 2009). Perfectly aligned reads represented miRNA loci and then imperfectly mapped reads (derived from isomiRs) were re-aligned to these loci using SHRiMP (Rumble et al., 2009). Aligned reads were quantified and normalized using reads per million mapped to miRNAs. Data are available from the Gene Expression Omnibus (GEO): accession number GSE114974.

### ChRO-seq read mapping

Read quality was assessed using FastQC. Adapters were trimmed from the 3’ ends of reads using cutadapt 1.16 (Martin, 2013) with a maximum 10% error rate, minimum 2 bp overlap, and minimum 20 quality score. Each read contained a 6 bp UMI enabling PCR deduplication by collapsing UMIs followed by UMI trimming using PRINSEQ lite 0.20.2 (Schmieder and Edwards, 2011). Processed reads with a minimum length of 15 bp were mapped to the hg38 genome modified with the addition of a single copy of the human Pol I ribosomal RNA complete repeating unit (GenBank U13369.1) with BWA 0.7.13 (Li and Durbin, 2010) using the BWA-backtrack algorithm. Each read was represented by a single base at the 5’ end of the read, corresponding to the 5’ end of the nascent RNA. Data was converted to bigwig format using bedtools 2.27.1 (Quinlan and Hall, 2010) and UCSC bedGraphToBigWig v4 (Kent et al., 2010) for visualization and identification of TREs. Bigwig files from identical conditions were merged and normalized to a total signal of 1×10^6^ prior to visualization.

### TRE identification

To identify TREs across all samples, bigwig files of the same strand from all samples (FLC and NML) were merged. This merged dataset was used to call TREs. TREs from all samples were identified with dREG (Danko et al., 2015; Wang et al., 2019) using the peak calling algorithm. Read counts were quantified within each TRE locus using the R package bigwig (https://github.com/andrelmartins/bigWig). Total read counts on the sense and antisense strands within each TRE across all samples were then imported into DESeq2 1.22.2 (Love et al., 2014). Analysis of TRE counts from ChRO-seq revealed they followed a negative binomial distribution similar to RNA-seq counts. Therefore, differential transcription analysis of TREs was performed with DESeq2 to identify TREs that were significantly differentially transcribed in FLC or NML.

### Differential gene transcription analysis

Gene definitions were obtained from GENCODE v25 annotations. To avoid counting reads from the paused polymerase peak, ChRO-seq signal was quantified on the sense strand from 500 bp downstream of the gene start until the annotated end of the gene. Genes were eliminated from the analysis if they were shorter than 1000 bp and if they were not protein coding, pseudogene, lincRNA, antisense, or miRNA genes. Like TRE counts, gene body counts from ChRO-seq followed a negative binomial distribution. Therefore, differential transcription analysis of genes was performed using DESeq2.

### TRE analyses

TRE annotation was performed using the annotatePeaks.pl function from HOMER (Heinz et al., 2010) based on GENCODE v25 annotations. Transcription factor motif enrichment analysis was performed using the findMotifsGenome.pl function from HOMER using “given” as the size parameter. The input (and background) peaks run were FLC-specific TREs (all TREs that were not identified as FLC-specific as background), NML-specific TREs (all TREs not identified as NML-specific), FLC-specific TREs without FOSL2/JUN or CREB motifs (all TREs not identified as FLC-specific), FLC-specific promoters (all promoters not identified as FLC-specific), and FLC-specific enhancers (all enhancers not identified as FLC-specific). CpG island annotations were downloaded from the UCSC Table Browser.

Hierarchical clustering was performed following DESeq2 normalization of read counts quantified from each TRE or gene body as described above. Clustering was performed in a pairwise manner using a correlation-based distance metric (1 - Spearman’s rho) using Ward’s minimum variance method. Proximal (−1000 to +100 bp from TSS) and distal TREs (the remaining TREs) were classified based on GENCODE v25 transcript annotations. Principal components analysis was performed following Variance Stabilizing Transformation from DESeq2.

TRE functional enrichment analysis was performed using GREAT (McLean et al., 2010). Briefly, TREs were converted from hg38 into hg19 coordinates using the UCSC liftOver tool. GREAT was run using default parameters (whole genome background, basal plus extension association rule). Gene ontology analyses, including KEGG 2016, ARCHS4, and PPI hub enrichment, were performed using Enrichr (Chen et al., 2013). Windows for enhancer density and gene-enhancer correlations were defined using gene coordinates based on GENCODE v25 annotations.

### FLC-specific enhancer hotspots

FLC-specific enhancer hotspots were identified using a method analogous to those previously described for super enhancers (Lovén et al., 2013; Whyte et al., 2013). First, distal TREs (TREs not overlapping -1000 to +100 bp from any TSS based on GENCODE v25 transcript annotations) were stitched together using a stitching distance of 12.5 kb. Read counts normalized by DESeq2 within each distal TRE were quantified in all samples, averaged for each TRE, and summed for each stitched enhancer. Stitched enhancers were ranked based on cumulative signal and a threshold for super enhancer identification was determined by drawing a line tangent to the signal curve. Stitched enhancers with more signal than the point identified by the tangent line were classified as FLC-specific enhancer hotspots.

Coordinates of known super enhancers were downloaded from SEdb (Jiang et al., 2019; http://www.licpathway.net/sedb/). ENCODE enhancer coordinates were downloaded from Search Candidate cis-Regulatory Elements by ENCODE (SCREEN, https://screen.wenglab.org/). FANTOM5 enhancer coordinates were downloaded from FANTOM5 (http://fantom.gsc.riken.jp/5/). NIH Roadmap Epigenomics enhancer coordinates were downloaded from Reg2Map: HoneyBadger (https://personal.broadinstitute.org/meuleman/reg2map/HoneyBadger_release/). FANTOM5 data was downloaded in hg38 coordinates. SEdb, ENCODE, and NIH Roadmap data were available and downloaded in hg19 coordinates and converted to hg38 coordinates using the UCSC liftOver tool.

### Gene-enhancer correlations

Transcriptional signal was quantified from enhancers and gene bodies as described above. Gene body and enhancer counts were normalized separately using DESeq2 and the Pearson correlation coefficients between the log_2_(normalized counts + 1) for genes and enhancers were calculated. To determine the statistical significance of the calculated correlation coefficients, we constructed a null distribution that consisted of correlation coefficients from genes and enhancers on different chromosomes. For gene-enhancer pairs within 100 kb of each other, we calculated the empirical p-value based on the null distribution and adjusted for multiple testing using the Benjamini-Hochberg (FDR) procedure (Benjamini and Hochberg, 1995).

### Visualization of RNA polymerase signal

Heatmaps and line graphs visualizing RNA polymerase signal were generated using deepTools 3.0.2 (Ramírez et al., 2016). Genomic loci snapshots were generated using Gviz 1.26.5 (Hahne and Ivanek, 2016).

### RNA-seq analysis

Read quality was assessed using FastQC. Reads were mapped to the hg38 genome with STAR 2.4.2a (Dobin et al., 2012). Transcripts were quantified with Salmon 0.8.2 (Patro et al., 2017) using GENCODE v25 transcript annotations. Normalization was performed using DESeq2. TCGA RNA-seq data were downloaded using TCGA-assembler 2 (Wei et al., 2018) as normalized counts. TCGA RNA-seq data for lncRNAs was downloaded from TANRIC (Li et al., 2015) as normalized counts. Cholangiocarcinoma RNA-seq data (Nakamura et al., 2015) was downloaded from the European Genome-Phenome Archive (accession number EGAS00001000950). GTEx Release V7 data was downloaded from the GTEx Portal as gene read counts and normalized with RNA-seq data from primary FLCs using DESeq2. Principal components analysis was performed using the most variable 1000 genes following variance stabilizing transformation (DESeq2).

### smRNA-seq analysis

Read quality was assessed using FastQC. Reads were trimmed, mapped, and quantified to the hg19 genome using miRquant 2.0, our previously described smRNA-seq analysis pipeline (Kanke et al., 2016). Briefly, reads were trimmed using Cutadapt and reads were mapped to the genome using Bowtie (Langmead et al., 2009). Perfectly aligned reads represented miRNA loci and imperfectly mapped reads (from isomiRs) were re-aligned to these loci using SHRiMP (Rumble et al., 2009). Aligned reads were quantified and normalized using reads per million mapped to miRNAs (RPMMM).

### Quantitative PCR

Total RNA was isolated using the Total RNA Purification Kit (Norgen Biotek) per manufacturer’s instructions. Reverse transcription was performed using the High Capacity RNA-to-cDNA Kit (Thermo Fisher Scientific). Gene expression was quantified with TaqMan Gene Expression Assays (Thermo Fisher Scientific) on a CFX96 Touch Real-Time System (Bio-Rad). RNA expression levels were normalized to *RPS9*. The TaqMan assays used are provided in Table S8.

### Western blotting

Protein lysates were prepared using RIPA buffer (Sigma, St. Louis, MO) supplemented with protease inhibitor cocktail (Sigma), phosphatase inhibitor cocktails 1 and 2 (Sigma), 1 mmol/L phenylmethylsulfonyl fluoride, 0.1% β-mercaptoethanol, and 1 mmol/L dithiothreitol. Protein concentration was measured using the Pierce BCA Protein Assay (Thermo Fisher Scientific) according to the manufacturer’s protocol. Lysates were subjected to sodium dodecyl sulfate polyacrylamide gel electrophoresis (SDS-PAGE) using 25 μg of lysate per lane under denaturing conditions in NuPAGE 10% Bis-Tris (Thermo Fisher Scientific) or homemade 12% Bis-Tris gels and transferred to PVDF membranes using a standard wet transfer protocol. Membranes were blocked with 5% dry nonfat milk in TBST and were probed with antibodies. Enhanced chemiluminescence reagent (GE Healthcare, Chicago, IL) was used for detection.

### siRNA transfection

AML12 cells were reverse transfected using Lipofectamine RNAiMAX (Thermo Fisher Scientific) per manufacturer’s instructions. Briefly, transfection mixes were assembled in wells by adding 1 pmol of the appropriate siRNA and 0.3 μL RNAiMAX (96-well plates) or 5 pmol siRNA and 1.5 μL RNAiMAX (6-well plates) according to manufacturer’s protocols. OptiMEM was added to each well and the plates were incubated for 15 minutes at room temperature. 1,500 cells (96-well plates) or 5,000 cells (6-well plates) were plated on top of transfection mixes. Cells were incubated for 24 hours prior to further treatment.

### Drug treatments

For AML12 cells seeded in 96-well plates, a series of ½-log unit dilutions of cobimetinib (10 mM stock in DMSO) were made in DMSO at 1000X final desired concentrations. From these stocks, 1:200 dilutions were made in fresh AML12 media. 30 μl of these starting dilutions was added to appropriate wells using a multi-channel pipettor. This results in a further 1:5 dilution and a final 1:1000 dilution with a final volume of 150 μl per well. Outer plate wells were filled with media and a no-cells/no-treatment set of wells was included for background. Cell plates were grown for a further 4-5 days.

FLC cells were seeded in 96-well plates at a density of 3200 cells/well in a volume of 100 μl per well for CellTiter-Glo (Promega, Madison, WI) assays. The following day, 2X concentrations of drug or DMSO were prepared. 50 μl of media was removed from each well and 50 μl of 2X drug or vehicle (final concentration 1X) was added to each well. Cells were incubated for 48 hours before assessing cell viability.

### Cell viability

Cell viability was assessed by CellTiter-Glo or crystal violet staining. For CellTiter-Glo, 96-well plates were removed from incubator and placed at room temperature for 30 minutes to equilibrate. Room temperature CellTiter-Glo reagent was added and cells were shaken for 2 minutes. Plates were then incubated for 10 min at room temperature. Luminescence was measured using a POLARStar Omega plate reader (BMG LabTech, Ortenberg, Germany; Em Filter – empty; Gain = 3600, orbital averaging ON, diameter = 5, cycles = 6) or a Synergy 2 Microplate Reader (Biotek, Winooski, VT; area scan; integration time = 0.50 seconds).

For crystal violet staining, AML12 cells were rinsed in PBS and fixed in 4% paraformaldehyde in PBS for 20 minutes. Two water washes were performed and cells were stained with 0.25% crystal violet in 10% methanol for 20 minutes. Finally, three water washes were performed and plates were allowed to dry at room temperature for at least 24 hours. Images were captured with a custom digital photography set-up on a Canon 5D, MkI with a Sigma 150-600 mm lens. To quantify crystal violet staining, dye was dissolved in 10% acetic acid (300 μl per well). An aliquot was removed to a clear 96-well plate and A590 absorbance was measured using a POLARStar Omega plate reader (BMG LabTech). Signal was kept in the linear range by 1:2 – 1:4 dilution with 10% acetic acid where necessary.

### Combination Index

Relative quantification values (RQVs) were calculated by normalizing the effect of drug treatment against DMSO controls following CellTiterGlo cell viability experiments. RQVs were generated from 4-12 replicates and were entered as effect values at appropriate drug dosages in single and combination drug treatments in CompuSyn Software using non-constant ratio design parameters. Individual Combination Index values are reported for each dose combination.

## Quantification and Statistical Analysis

### Statistics

Statistical analyses were performed using R (3.5.0). Statistical significance was primarily determined by two-sided Welch’s t-tests or Mann-Whitney U tests as indicated in the figure legends. All alternative statistical tests that were used are noted in the text or figure legends. Statistical comparisons between dose response curves were performed using the R package *drc* (3.0-1) by fitting log-logistic models (with the LL.4 function) to the data. Two models were fit to the data (one with and one without siRNA) and were compared using a F-test (with the *anova* function) to determine statistical significance. P<0.05 was considered statistically significant unless otherwise noted. *p<0.05, **p<0.01, ***p<0.001.

### Data and Code Availability

ChRO-seq and RNA-seq data are currently being deposited into the European Genome-Phenome Archive (EGA). An accession number to the dataset will be provided as soon as it becomes available. The code used in this study is available from the corresponding author upon request.

## End Matter

### Author Contributions and Notes

Conceptualization, T.A.D. and P.S.; Methodology, T.A.D., R.S., A.B.F., F.D.S, R.K.M., R.P.B., and P.S.; Software, T.A.D. and M.K.; Analysis, T.A.D., R.S., A.B.F., F.D.S, R.K.M., R.P.B., M.K., and P.S.; Investigation, T.A.D., R.S., A.B.F., F.D.S, R.K.M., R.P.B., and A.P.M.; Resources, C.G.D. and J.D.S.; Writing – Original Draft, T.A.D. and P.S.; Writing – Review & Editing, all authors, especially T.A.D., R.S., F.D.S., J.D.S., and P.S.; Visualization, T.A.D. and F.D.S.; Supervision, P.S.

The authors declare no conflict of interest.

## Acknowledgments

The authors would like to thank members of the Sethupathy laboratory for helpful discussions regarding this study; Tinyi Chu and Ed Rice for discussions and assistance in experimental protocols and computational analysis of ChRO-seq; and Selina Ruzi for assistance in drawing Figure 1A. The authors also thank John Hopper, Dr. Mark Furth, Tom Stockwell, Marna Davis, and the entire Fibrolamellar Cancer Foundation for their support of the patients and the research network. We thank Ilya Finkelstein for the bioRxiv template.

## References

Benjamini, Y., and Hochberg, Y. (1995). Controlling the False Discovery Rate : A Practical and Powerful Approach to Multiple Testing Author. J. R. Stat. Soc. 57, 289–300.

Blackwood, E.M., and Kadonaga, J.T. (1998). Going the Distance: A Current View of Enhancer Action. Science 281, 60–63.

Boyd, N.H., Walker, K., Fried, J., Hackney, J.R., McDonald, P.C., Benavides, G.A., Spina, R., Audia, A., Scott, S.E., Libby, C.J., et al. (2017). Addition of carbonic anhydrase 9 inhibitor SLC-0111 to temozolomide treatment delays glioblastoma growth in vivo. JCI Insight 2, e92928.

Chen, E.Y., Tan, C.M., Kou, Y., Duan, Q., Wang, Z., Meirelles, G.V., Clark, N.R., and Ma’ayan, A. (2013). Enrichr: interactive and collaborative HTML5 gene list enrichment analysis tool. BMC Bioinformatics 14, 128.

Chen, Z., Li, J., Lin, S., Cao, C., Gimbrone, N.T., Yang, R., Fu, D.A., Carper, M.B., Haura, E.B., Schabath, M.B., et al. (2016). cAMP / CREB-regulated LINC00473 marks LKB1-inactivated lung cancer and mediates tumor growth. J. Clin. Invest. 126, 2267–2279.

Chen, Z., Lin, S., Li, J.-L., Ni, W., Guo, R., Lu, J., Kaye, F.J., and Wu, L. (2018). CRTC1-MAML2 fusion-induced lncRNA LINC00473 expression maintains the growth and survival of human mucoepidermoid carcinoma cells. Oncogene 37, 1885–1895.

Chu, T., Rice, E., Booth, G., Salamanca, H., Wang, Z., Core, L., Longo, S., Corona, R., Chin, L., Lis, J., et al. (2018). Chromatin run-on and sequencing maps the transcriptional regulatory landscape of glioblastoma multiforme. Nat. Genet. 50, 1553–1564.

Corces, M.R., Buenrostro, J.D., Wu, B., Greenside, P.G., Chan, S.M., Koenig, J.L., Snyder, M.P., Pritchard, J.K., Kundaje, A., Greenleaf, W.J., et al. (2016). Lineage-specific and single-cell chromatin accessibility charts human hematopoiesis and leukemia evolution. Nat. Genet. 48, 1193–1203.

Corces, M.R., Granja, J.M., Shams, S., Louie, B.H., Seoane, J.A., Zhou, W., Silva, T.C., Groeneveld, C., Wong, C.K., Cho, W., et al. (2018). The chromatin accessibility landscape of primary human cancers. Science 362, eaav1898.

Core, L., Waterfall, J., and Lis, J. (2008). Nascent RNA sequencing reveals widespread pausing and divergent initiation at human promoters. Science 322, 1845–1848.

Cornella, H., Alsinet, C., Sayols, S., Zhang, Z., Hao, K., Cabellos, L., Hoshida, Y., Villanueva, A., Thung, S., Ward, S.C., et al. (2014). Unique genomic profile of fibrolamellar hepatocellular carcinoma. Gastroenterology 148, 806–818.

Craig, J.R., Peters, R.L., Edmondson, H.A., and Omata, M. (1980). Fibrolamellar carcinoma of the liver: a tumor of adolescents and young adults with distinctive clinico-pathologic features. Cancer 46, 372–379.

Danko, C.G., Hyland, S.L., Core, L.J., Martins, A.L., Waters, C.T., Lee, H.W., Cheung, V.G., Kraus, W.L., Lis, J.T., and Siepel, A. (2015). Identification of active transcriptional regulatory elements from GRO-seq data. Nat. Methods 12, 433–438.

Dinh, T.A., Vitucci, E.C.M., Wauthier, E., Graham, R.P., Pitman, W.A., Oikawa, T., Chen, M., Silva, G.O., Greene, K.G., Torbenson, M.S., et al. (2017). Comprehensive analysis of The Cancer Genome Atlas reveals a unique gene and non-coding RNA signature of fibrolamellar carcinoma. Sci. Rep. 7, 44653.

Dinh, T.A., Jewell, M.L., Kanke, M., Francisco, A., Sritharan, R., Turnham, R.E., Lee, S., Kastenhuber, E.R., Wauthier, E., Guy, C.D., et al. (2019). MicroRNA-375 Suppresses the Growth and Invasion of Fibrolamellar Carcinoma. Cell. Mol. Gastroenterol. Hepatol. 7, 803–817.

Dobin, A., Davis, C.A., Schlesinger, F., Drenkow, J., Zaleski, C., Jha, S., Batut, P., Chaisson, M., and Gingeras, T.R. (2012). STAR: ultrafast universal RNA-seq aligner. Bioinformatics 29, 15–21.

Doyen, J., Parks, S.K., Marcié, S., Pouysségur, J., and Chiche, J. (2013). Knock-down of hypoxia-induced carbonic anhydrases IX and XII radiosensitizes tumor cells by increasing intracellular acidosis. Front. Oncol. 2, 199.

Dunham, I., Kundaje, A., Aldred, S.F., Collins, P.J., Davis, C.A., Doyle, F., Epstein, C.B., Frietze, S., Harrow, J., Kaul, R., et al. (2012). An integrated encyclopedia of DNA elements in the human genome. Nature 489, 57–74.

Engelholm, L.H., Riaz, A., Serra, D., Dagnæs-Hansen, F., Johansen, J. V., Santoni-Rugiu, E., Hansen, S.H., Niola, F., and Frödin, M. (2017). CRISPR/Cas9 engineering of adult mouse liver demonstrates that the Dnajb1–Prkaca gene fusion is sufficient to induce tumors resembling fibrolamellar hepatocellular carcinoma. Gastroenterology 153, 1662–1673.

Farber, B.A., Lalazar, G., Simon, E.P., Hammond, W.J., Requena, D., Bhanot, U.K., La Quaglia, M.P., and Simon, S.M. (2018). Non coding RNA analysis in fibrolamellar hepatocellular carcinoma. Oncotarget 9, 10211–10227.

Filippakopoulos, P., Qi, J., Picaud, S., Shen, Y., Smith, W.B., Fedorov, O., Morse, E.M., Keates, T., Hickman, T.T., Felletar, I., et al. (2010). Selective inhibition of BET bromodomains. Nature 468, 1067–1073.

Franco, H.L., Nagari, A., Malladi, V.S., Li, W., Xi, Y., Richardson, D., Allton, K.L., Tanaka, K., Li, J., Murakami, S., et al. (2018). Enhancer transcription reveals subtype-specific gene expression programs controlling breast cancer pathogenesis. Genome Res. 28, 159–170.

Ginty, D.D., Bonni, A., and Greenberg, M.E. (1994). Nerge growth factor activates a Ras-dependent protein kinase that stimulates c-fos transcription via phosphorlation of CREB. Cell 77, 713–725.

Graham, R.P., Jin, L., Knutson, D.L., Kloft-Nelson, S.M., Greipp, P.T., Waldburger, N., Roessler, S., Longerich, T., Roberts, L.R., Oliveira, A.M., et al. (2015). DNAJB1-PRKACA is specific for fibrolamellar carcinoma. Mod. Pathol. 28, 822–829.

Griffith, O.L., M. Griffith, K. Krysiak, V. Magrini, A. Ramu, Z.L. Skidmore, J. Kunisaki, R. Austin, S. McGrath, J. Zhang, et al. (2016). A genomic case study of mixed fibrolamellar hepatocellular carcinoma. Ann. Oncol. 27, 1148–1154.

Van Groningen, T., Koster, J., Valentijn, L.J., Zwijnenburg, D.A., Akogul, N., Hasselt, N.E., Broekmans, M., Haneveld, F., Nowakowska, N.E., Bras, J., et al. (2017). Neuroblastoma is composed of two super-enhancer-associated differentiation states. Nat. Genet. 49, 1261–1266.

Gryder, B.E., Yohe, M.E., Chou, H.C., Zhang, X., Marques, J., Wachtel, M., Schaefer, B., Sen, N., Song, Y., Gualtieri, A., et al. (2017). PAX3-FOXO1 establishes myogenic super enhancers and confers BET bromodomain vulnerability. Cancer Discov. 7, 884–899.

Hahne, F., and Ivanek, R. (2016). Visualizing Genomic Data Using Gviz and Bioconductor. In Statistical Genomics: Methods and Protocols, E. Mathé, and S. Davis, eds. (New York, NY: Springer New York), pp. 335–351.

Han, P.-B., Ji, X.-J., Zhang, M., and Gao, L.-Y. (2018). Upregulation of lncRNA LINC00473 promotes radioresistance of HNSCC cells through activating Wnt/beta-catenin signaling pathway. Eur. Rev. Med. Pharmacol. Sci. 22, 7305–7313.

Heintzman, N.D., Hon, G.C., Hawkins, R.D., Kheradpour, P., Stark, A., Harp, L.F., Ye, Z., Lee, L.K., Stuart, R.K., Ching, C.W., et al. (2009). Histone modifications at human enhancers reflect global cell-type-specific gene expression. Nature 459, 108–112.

Heinz, S., Benner, C., Spann, N., Bertolino, E., Lin, Y.C., Laslo, P., Cheng, J.X., Murre, C., Singh, H., and Glass, C.K. (2010). Simple combinations of lineage-determining transcription factors prime cis-regulatory elements required for macrophage and B cell identities. Mol. Cell 38, 576–589.

Hnisz, D., Abraham, B.J., Lee, T.I., Lau, A., Saint-André, V., Sigova, A.A., Hoke, H.A., and Young, R.A. (2013). Super-enhancers in the control of cell identity and disease. Cell 155, 934–947.

Honeyman, J.N., Simon, E.P., Robine, N., Chiaroni-Clarke, R., Darcy, D.G., Lim, I.I.P., Gleason, C.E., Murphy, J.M., Rosenberg, B.R., Teegan, L., et al. (2014). Detection of a recurrent DNAJB1-PRKACA chimeric transcript in fibrolamellar hepatocellular carcinoma. Science 343, 1010–1014.

Hsieh, M., Chen, K., Chiou, H., and Hsieh, Y. (2010). Carbonic anhydrase XII promotes invasion and migration ability of MDA-MB-231 breast cancer cells through the p38 MAPK signaling pathway. Eur. J. Cell Biol. 89, 598–606.

Januchowski, R., Zawierucha, P., Rucinski, M., Andrzejewska, M., Wojtowicz, K., Nowicki, M., and Zabel, M. (2014). Drug transporter expression profiling in chemoresistant variants of the A2780 ovarian cancer cell line. Biomed. Pharmacother. 68, 447–453.

Javierre, B.M., Burren, O.S., Wilder, S.P., Kreuzhuber, R., Hill, S.M., Sewitz, S., Cairns, J., Wingett, S.W., Várnai, C., Thiecke, M.J., et al. (2016). Lineage-specific genome architecture links enhancers and non-coding disease variants to target gene promoters. Cell 167, 1369–1384.

Jiang, Y., Qian, F., Bai, X., Liu, Y., Wang, Q., Ai, B., Han, X., Shi, S., Zhang, J., Li, X., et al. (2019). SEdb: A comprehensive human super-enhancer database. Nucleic Acids Res. 47, D235–D243.

Kanke, M., Baran-Gale, J., Villanueva, J., and Sethupathy, P. (2016). miRquant 2.0 : an expanded tool for accurate annotation and quantification of microRNAs and their isomiRs from small RNA-sequencing data. J. Integr. Bioinform. 13, 307.

Karin, M. (1995). The regulation of AP-1 activity by mitogen-activates protein kinases. J. Biol. Chem. 270, 16483–16486.

Kastenhuber, E.R., Lalazar, G., Tschaharganeh, D.F., Houlihan, S.L., Baslan, T., Chen, C.-C., Requena, D., Tian, S., Bosbach, B., Wilkinson, J.E., et al. (2017). DNAJB1-PRKACA fusion kinase drives tumorigenesis and interacts with β-catenin and the liver regenerative response. Proc. Natl. Acad. Sci. 114, 13076–13084.

Keller, D.M., Clark, E.A., and Goodman, R.H. (2012). Regulation of microRNA-375 by cAMP in Pancreatic β-Cells. Mol. Endocrinol. 26, 989–999.

Kent, W.J., Zweig, A.S., Barber, G., Hinrichs, A.S., and Karolchik, D. (2010). BigWig and BigBed: Enabling browsing of large distributed datasets. Bioinformatics 26, 2204–2207.

Kim, T.-K., Hemberg, M., Gray, J.M., Costa, A.M., Bear, D.M., Wu, J., Harmin, D.A., Laptewicz, M., Barbara-Haley, K., Kuersten, S., et al. (2010). Widespread transcription at neuronal activity-regulated enhancers. Nature 465, 182–187.

Kopecka, J., Rankin, G.M., Salaroglio, I.C., Poulsen, S.-A., and Riganti, C. (2016). P-glycoprotein-mediated chemoresistance is reversed by carbonic anhydrase XII inhibitors. Oncotarget 7, 85861–85875.

Kwak, H., Fuda, N.J., Core, L.J., and Lis, J.T. (2013). Precise maps of RNA polymerase reveal how promoters direct initiation and pausing. Science 339, 950–953.

Lachmann, A., Torre, D., Keenan, A.B., Jagodnik, K.M., Lee, H.J., Wang, L., Silverstein, M.C., and Ma’ayan, A. (2018). Massive mining of publicly available RNA-seq data from human and mouse. Nat. Commun. 9, 1366.

Langmead, B., Trapnell, C., Pop, M., and Salzberg, S.L. (2009). Ultrafast and memory-efficient alignment of short DNA sequences to the human genome. Genome Biol. 10, R25.

Lee, J.H., Choi, S.I., Kim, R.K., Cho, E.W., and Kim, I.G. (2018). Tescalcin/c-Src/IGF1Rβ-mediated STAT3 activation enhances cancer stemness and radioresistant properties through ALDH1. Sci. Rep. 8, 1–13.

Li, H., and Durbin, R. (2010). Fast and accurate long-read alignment with Burrows-Wheeler transform. Bioinformatics 26, 589–595.

Li, C., Singh, B., Graves-Deal, R., Ma, H., Starchenko, A., Fry, W.H., Lu, Y., Wang, Y., Bogatcheva, G., Khan, M.P., et al. (2017). Three-dimensional culture system identifies a new mode of cetuximab resistance and disease-relevant genes in colorectal cancer. Proc. Natl. Acad. Sci. 114, E2852–E2861.

Li, J., Han, L., Roebuck, P., Diao, L., Liu, L., Yuan, Y., Weinstein, J.N., and Liang, H. (2015). TANRIC: an interactive open platform to explore the function of lncRNAs in cancer. Cancer Res. 75, 3728–3737.

Liu, X., Krawczyk, E., Suprynowicz, F.A., Palechor-Ceron, N., Yuan, H., Dakic, A., Simic, V., Zheng, Y., Sripadhan, P., Chen, C., et al. (2017). Conditional reprogramming and long-term expansion of normal and tumor cells from human biospecimens. Nat. Protoc. 12, 439–451.

Long, H.K., Prescott, S.L., and Wysocka, J. (2016). Ever-changing landscapes: transcriptional enhancers in development and evolution. Cell 167, 1170–1187.

Love, M.I., Huber, W., and Anders, S. (2014). Moderated estimation of fold change and dispersion for RNA-seq data with DESeq2. Genome Biol. 15, 550.

Lovén, J., Hoke, H.A., Lin, C.Y., Lau, A., Orlando, D.A., Vakoc, C.R., Bradner, J.E., Lee, T.I., and Young, R.A. (2013). Selective inhibition of tumor oncogenes by disruption of super-enhancers. Cell 153, 320–334.

Ma, T.C., Barco, A., Ratan, R.R., and Willis, D.E. (2014). CAMP-responsive element-binding protein (CREB) and cAMP co-regulate activator protein 1 (AP1)-dependent regeneration-associated gene expression and neurite growth. J. Biol. Chem. 289, 32914–32925.

Mack, S.C., Pajtler, K.W., Chavez, L., Okonechnikov, K., Bertrand, K.C., Wang, X., Erkek, S., Federation, A., Song, A., Lee, C., et al. (2018). Therapeutic targeting of ependymoma as informed by oncogenic enhancer profiling. Nature 553, 101–105.

Mahat, D.B., Kwak, H., Booth, G.T., Jonkers, I.H., Danko, C.G., Patel, R.K., Waters, C.T., Munson, K., Core, L.J., and Lis, J.T. (2016). Base-pair-resolution genome-wide mapping of active RNA polymerases using precision nuclear run-on (PRO-seq). Nat. Protoc. 11, 1455–1476.

Malouf, G.G., Job, S., Paradis, V., Fabre, M., Brugières, L., Saintigny, P., Vescovo, L., Belghiti, J., Branchereau, S., Faivre, S., et al. (2014). Transcriptional profiling of pure fibrolamellar hepatocellular carcinoma reveals an endocrine signature. Hepatology 59, 2228–2237.

Man, C.H., Lam, S.S.Y., Sun, M.K.H., Chow, H.C.H., Gill, H., Kwong, Y.L., and Leung, A.Y.H. (2014). A novel tescalcin-sodium/hydrogen exchange axis underlying sorafenib resistance in FLT3-ITD+ AML. Blood 123, 2530–2539.

Maniaci, V., Davidson, B.R., Rolles, K., Dhillon, A.P., Hackshaw, A., Begent, R.H., and Meyer, T. (2009). Fibrolamellar hepatocellular carcinoma - Prolonged survival with multimodality therapy. Eur. J. Surg. Oncol. 35, 617–621.

Martin, M. (2013). Cutadapt removes adapter sequences from high-throughput sequencing reads. EMBnet.Journal 17, 10–12.

McLean, C.Y., Bristor, D., Hiller, M., Clarke, S.L., Schaar, B.T., Lowe, C.B., Wenger, A.M., and Bejerano, G. (2010). GREAT improves functional interpretation of cis-regulatory regions. Nat. Biotechnol. 28, 495–501.

Nakamura, H., Arai, Y., Totoki, Y., Shirota, T., Elzawahry, A., Kato, M., Hama, N., Hosoda, F., Urushidate, T., Ohashi, S., et al. (2015). Genomic spectra of biliary tract cancer. Nat. Genet. 47, 1003–1010.

Nord, A.S., Blow, M.J., Attanasio, C., Akiyama, J.A., Holt, A., Hosseini, R., Phouanenavong, S., Plajzer-Frick, I., Shoukry, M., Afzal, V., et al. (2013). Rapid and pervasive changes in genome-wide enhancer usage during mammalian development. Cell 155, 1521–1531.

Oikawa, T., Wauthier, E., Dinh, T.A., Selitsky, S.R., Reyna-Neyra, A., Carpino, G., Levine, R., Cardinale, V., Klimstra, D., Gaudio, E., et al. (2015). Model of fibrolamellar hepatocellular carcinomas reveals striking enrichment in cancer stem cells. Nat. Commun. 6, 8070.

Ooi, W.F., Xing, M., Xu, C., Yao, X., Ramlee, M.K., Lim, M.C., Cao, F., Lim, K., Babu, D., Poon, L.F., et al. (2016). Epigenomic profiling of primary gastric adenocarcinoma reveals super-enhancer heterogeneity. Nat. Commun. 7, 1–17.

Parker, S.C.J., Stitzel, M.L., Taylor, D.L., Orozco, J.M., Erdos, M.R., Akiyama, J.A., van Bueren, K.L., Chines, P.S., Narisu, N., Program, N.C.S., et al. (2013). Chromatin stretch enhancer states drive cell-specific gene regulation and harbor human disease risk variants. Proc. Natl. Acad. Sci. 110, 17921–17926.

Patro, R., Duggal, G., Love, M.I., Irizarry, R.A., and Kingsford, C. (2017). Salmon provides fast and bias-aware quantification of transcript expression. Nat. Methods 14, 417–419.

Peeters, J.G.C., Vervoort, S.J., Tan, S.C., Mijnheer, G., de Roock, S., Vastert, S.J., Nieuwenhuis, E.E.S., van Wijk, F., Prakken, B.J., Creyghton, M.P., et al. (2015). Inhibition of Super-Enhancer Activity in Autoinflammatory Site-Derived T Cells Reduces Disease-Associated Gene Expression. Cell Rep. 12, 1986–1996.

Pott, S., and Lieb, J.D. (2015). What are super-enhancers? Nat. Genet. 47, 8–12.

Quinlan, A.R., and Hall, I.M. (2010). BEDTools: A flexible suite of utilities for comparing genomic features. Bioinformatics 26, 841–842.

Ramírez, F., Ryan, D.P., Grüning, B., Bhardwaj, V., Kilpert, F., Richter, A.S., Heyne, S., Dündar, F., and Manke, T. (2016). deepTools2: a next generation web server for deep-sequencing data analysis. Nucleic Acids Res. 44, W160–W165.

Reitmair, A., Sachs, G., Im, W.B., and Wheeler, L. (2012). C6orf176: a novel possible regulator of cAMP-mediated gene expression. Physiol. Genomics 44, 152–161.

Rumble, S.M., Lacroute, P., Dalca, A. V, Fiume, M., Sidow, A., and Brudno, M. (2009). SHRiMP: Accurate Mapping of Short Color-space Reads. PLoS Comput. Biol. 5, e1000386.

De Santa, F., Barozzi, I., Mietton, F., Ghisletti, S., Polletti, S., Tusi, B.K., Muller, H., Ragoussis, J., Wei, C.-L., and Natoli, G. (2010). A large fraction of extragenic RNA Pol II transcription sites overlap enhancers. PLoS Biol. 8, e1000384.

Sanyal, S., Sandstrom, D.J., Hoeffer, C.A., and Ramaswami, M. (2002). AP-1 functions upstream of creb to control synaptic plasticity in drosophila. Nature 416, 870–874.

Schmieder, R., and Edwards, R. (2011). Quality control and preprocessing of metagenomic datasets. Bioinformatics 27, 863–864.

Shaywitz, A.J., and Greenberg, M.E. (1999). CREB: A Stimulus-Induced Transcription Factor Activated by a Diverse Array of Extracellular Signals. Annu. Rev. Biochem. 68, 821–861.

Shi, C., Yang, Y., Yu, J., Meng, F., Zhang, T., and Gao, Y. (2017). The long noncoding RNA LINC00473, a target of microRNA 34a, promotes tumorigenesis by inhibiting ILF2 degradation in cervical cancer. Am. J. Cancer Res. 7, 2157–2168.

Simon, E.P., Freije, C.A., Farber, B.A., Lalazar, G., Darcy, D.G., Honeyman, J.N., Chiaroni-Clarke, R., Dill, B.D., Molina, H., Bhanot, U.K., et al. (2015). Transcriptomic characterization of fibrolamellar hepatocellular carcinoma. Proc. Natl. Acad. Sci. 112, E5916–E5925.

Sorenson, E.C., Khanin, R., Bamboat, Z.M., Cavnar, M.J., Kim, T.S., Sadot, E., Zeng, S., Greer, J.B., Seifert, A.M., Cohen, N.A., et al. (2017). Genome and transcriptome profiling of fibrolamellar hepatocellular carcinoma demonstrates p53 and IGF2BP1 dysregulation. PLoS One 12, e0176562.

Stipa, F., Yoon, S.S., Liau, K.H., Fong, Y., Jarnagin, W.R., D’Angelica, M., Abou-Alfa, G., Blumgart, L.H., and DeMatteo, R.P. (2006). Outcome of patients with fibrolamellar hepatocellular carcinoma. Cancer 106, 1331–1338.

Torbenson, M. (2012). Fibrolamellar carcinoma: 2012 update. Scientifica 2012, 1–15.

Turnham, R.E., Smith, F.D., Kenerson, H.L., Omar, M.H., Golkowski, M., Garcia, I., Bauer, R., Lau, H.-T., Sullivan, K.M., Langeberg, L.K., et al. (2019). An acquired scaffolding function of the DNAJ-PKAc fusion contributes to oncogenic signaling in fibrolamellar carcinoma. Elife 8, e44187.

Wang, L., Zhang, X., Sheng, L., Qiu, C., and Luo, R. (2018). LINC00473 promotes the Taxol resistance via miR-15a in colorectal cancer. Biosci. Rep. 38, BSR20180790.

Wang, Z., Chu, T., Choate, L.A., and Danko, C.G. (2019). Identification of regulatory elements from nascent transcription using dREG. Genome Res. 29, 293–303.

Wei, L., Jin, Z., Yang, S., Xu, Y., Zhu, Y., and Ji, Y. (2018). TCGA-assembler 2: software pipeline for retrieval and processing of TCGA/CPTAC data. Bioinformatics 34, 1615–1617.

Whyte, W.A., Orlando, D.A., Hnisz, D., Abraham, B.J., Lin, C.Y., Kagey, M.H., Rahl, P.B., Lee, T.I., and Young, R.A. (2013). Master transcription factors and mediator establish super-enhancers at key cell identity genes. Cell 153, 307–319.

Wu, G.-Y., Deisseroth, K., and Tsien, R.W. (2001). Activity-dependent CREB phosphorylation: Convergence of a fast, sensitive calmodulin kinase pathway and a slow, less sensitive mitogen-activated protein kinase pathway. Proc. Natl. Acad. Sci. 98, 2808–2813.

Xing, J., Ginty, D.D., and Greenberg, M.E. (1996). Coupling of the RAS-MAPK pathway to gene activation by RSK2, a growth factor-regulated CREB kinase. Science 273, 959–963.

Xu, L., Hazard, F., Zmoos, A., Jahchan, N., Chaib, H., PM, G., Rangaswami, A., Snyder, M., and Sage, J. (2014). Genomic analysis of fibrolamellar hepatocellular carcinoma. Hum. Mol. Genet. 24, 1–42.

Yoo, C.W., Nam, B.-H., Kim, J.-Y., Shin, H.-J., Lim, H., Lee, S., Lee, S.-K., Lim, M.-C., and Song, Y.-J. (2010). Carbonic anhydrase XII expression is associated with histologic grade of cervical cancer and superior radiotherapy outcome. Radiat. Oncol. 5, 101.

